# Benchmarking methods for mapping functional connectivity in the brain

**DOI:** 10.1101/2024.05.07.593018

**Authors:** Zhen-Qi Liu, Andrea I. Luppi, Justine Y. Hansen, Ye Ella Tian, Andrew Zalesky, B.T. Thomas Yeo, Ben D. Fulcher, Bratislav Misic

**Affiliations:** Montréal Neurological Institute, McGill University, Montréal, Canada; Melbourne Neuropsychiatric Centre, The University of Melbourne, Melbourne, Australia; Yong Loo Lin School of Medicine, National University of Singapore, Singapore; School of Physics, The University of Sydney, Sydney, Australia

## Abstract

The networked architecture of the brain promotes synchrony among neuronal populations and the emergence of coherent dynamics. These communication patterns can be comprehensively mapped using noninvasive functional imaging, resulting in functional connectivity (FC) networks. Despite its popularity, FC is a statistical construct and its operational definition is arbitrary. While most studies use zero-lag Pearson’s correlations by default, there exist hundreds of pairwise interaction statistics in the broader scientific literature that can be used to estimate FC. How the organization of the FC matrix varies with the choice of pairwise statistic is a fundamental methodological question that affects all studies in this rapidly growing field. Here we comprehensively benchmark the topological and geometric organization, neurobiological associations, and cognitive-behavioral relevance of FC matrices computed using a large library of 239 pairwise interaction statistics. We comprehensively investigate how canonical features of FC networks vary with the choice of pairwise statistic, including (1) hub mapping, (2) weight-distance trade-offs, (3) structure–function coupling, (4) correspondence with other neurophysiological networks, (5) individual fingerprinting, and (6) brain–behavior prediction. We find substantial quantitative and qualitative variation across FC methods. Throughout, we observe that measures such as covariance (full correlation), precision (partial correlation) and distance display multiple desirable properties, including close correspondence with structural connectivity, the capacity to differentiate individuals and to predict individual differences in behavior. Using information flow decomposition, we find that differences among FC methods may arise from differential sensitivity to the underlying mechanisms of inter-regional communication, with some more sensitive to redundant and some to synergistic information flow. In summary, our report highlights the importance of tailoring a pairwise statistic to a specific neurophysiological mechanism and research question, providing a blueprint for future studies to optimize their choice of FC method.

## INTRODUCTION

The brain is a network of anatomically connected and perpetually interacting neuronal populations [1]. Its spectrum of functions—from perception to cognition to action—depends on inter-regional signaling [13]. Over the past 20 years, the dominant paradigm to infer inter-regional signaling has been to estimate functional connectivity (FC) [23, 24, 55, 64, 75, 89, 111, 148, 182, 207]. Regional time-series of metabolic, electromagnetic or haemodynamic neural activity are first recorded, and systematic co-activation between regions is then estimated and used to map FC networks [22, 39, 87, 153, 174, 176, 177, 206].

Perhaps the most widespread paradigm of estimating FC networks is using task-free or resting-state fMRI [22, 87, 178, 191, 204]. Neuronal population dynamics, in this setting, are recorded without task instruction or stimulation; the resulting “intrinsic” functional connectivity is thought to reflect spontaneous neural activity. Intrinsic functional patterns are highly organized [138, 210], reproducible [122], individual-specific [51, 114], correlated with structural connectivity [73, 183] and comparable to task-driven co-activation patterns [36, 175].

Unlike structural connectivity, which represents anatomical connections, functional connectivity is a statistical construct and does not represent a physical entity [87, 128–131]. As a result, there is no straightforward “ground truth” and how FC is estimated is a subjective methodological choice made by each individual researcher. Although multiple methods have been proposed, the most common method remains the simple zero-lag linear (Pearson’s) correlation coefficient. Yet, the broader scientific literature on estimating pairwise interactions among random variables is rich and vast, including those that capture nonlinear dependencies and time-lagged interactions [26, 32, 33]. How FC matrices vary with the choice of pairwise statistic is a fundamental methodological question that affects all studies in this field, limiting our understanding of the brain’s functional organization, as well as our capacity to develop optimized algorithms for structure–function coupling, individual fingerprinting and brain–behavior prediction [15, 27, 29, 35, 38, 50, 76, 102, 134, 148, 154, 157, 174, 176–178, 191, 198, 204, 208].

Here we comprehensively benchmark multiple features of resting-state FC using 239 pairwise interaction statistics. We first chart the similarities and differences among broad families of statistics. We then investigate how commonly studied features of the FC matrix— such as hubs, relationships with physical distance and structural connectivity—vary with the choice of pairwise statistic. We next show that individual differences in FC organization, including fingerprinting and brain– behavior relationships, also depend on choice of pairwise statistic. Finally, we use an information-theoretic decomposition to study how pairwise statistics capture different mechanisms of information flow.

## RESULTS

Pairwise statistics were derived for *N* = 326 unrelated healthy young adults from the Human Connectome Project (HCP) [193]. Functional time series were taken from HCP S1200 release minimal preprocessing pipeline with ICA-FIX cleaning [59, 155]. We used the *pyspi* package to estimate 239 pairwise statistics from 49 pairwise interaction measures in 6 families of statistics, yielding 239 FC matrices [32] for each participant. All main text results are shown for the undirected component of the matrices (upper triangular vector), and in the Schaefer100*×*7 atlas [161]. For other atlases and alternative processing choices, see the *Sensitivity Analysis* section.

### Massive profiling of pairwise interaction statistics

Figure 1 (top) shows the edge-wise similarities between the 239 FC matrices. Pairwise statistics are stratified according to the broad model family from which they are derived (e.g., information theoretic, spectral, etc.). The 49 pairwise measures are listed on the right, as well as the number of variants of each measure, which we refer to as pairwise statistics (239 total) [32].

**Figure 1.**
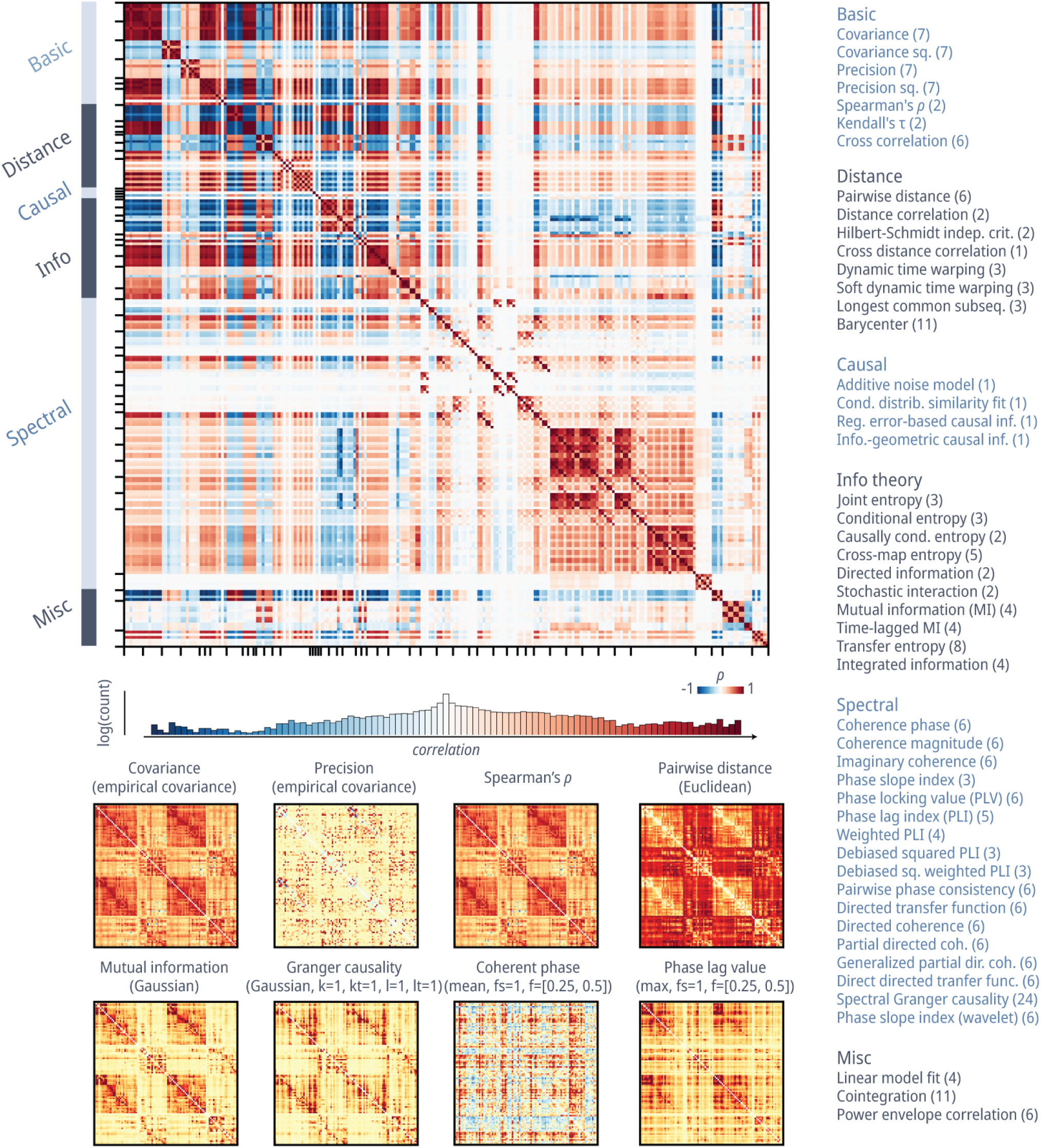
Massive profiling of pairwise interaction statistics for resting-state functional activity across the brain. Pairwise statistics for functional time series were estimated between all pairs of brain regions to generate 239 distinct FC matrices. **(Upper left)** Group-average similarity between all pairs of 239 pairwise statistics. Edge-wise similarities between individual pairwise statistics were quantified using Spearman’s rank correlation (*ρ*) for each participant, and then averaged across participants. The histogram of similarity values is shown below the matrix. The color represents [*−*1, 1], and bar height represents log-transformed count in each bin within the range of [*−*1, 1]. **(Lower left)** Group-average matrices for exemplar statistics calculated between pairs of time series. The annotation above each matrix denotes the broader family of the statistic and (in parentheses) details for the specific statistic. **(Right)** List of 239 pairwise statistics grouped into 49 measures across 6 major model families, following the categorization of Cliff et al. [32]. Numbers in parentheses indicate the number of specific variants of the statistics calculated for the measure. The colorbar covers only positive values (0th to 97.5th percentile; in red) for statistics with only positive values, and covers both negative values (0th percentile to zero; in blue) and positive values (zero to 97.5th percentile; in red) otherwise. A detailed list of the 239 pairwise statistics can be found in Table S1. The variance of the similarity matrix across subjects and runs can be found in Fig. S1.

Pairwise statistics are highly organized and form clusters that reflect families of statistics. For reference, the conventional zero-lag Pearson’s correlation is shown as the *covariance* family, and partial correlation is shown in the *precision* family in all figures. Some statistics are, by definition, highly similar to others. The most widely used family of statistics for FC calculation, *covariance* estimators, for example, are most correlated to *correlation, distance correlation*, and *mutual information* estimators. As expected, these measures of similarity tend to be highly anticorrelated with measures of dissimilarity such *precision, distance*, and *entropy*. Others—for example, spectral measures—show mild to moderate correlation with most other measures. Importantly, the correlations among the pairwise statistics distribute widely across the positive to negative range. This suggests that different method used to compute the FC matrix may yield networks with very different configurations. Indeed, Fig. 1 (bottom) shows eight sample FC matrices. The matrices are visually different: for example, some show clear block-like structure, while others do not.

### Benchmarking topological and geometric organization

If pairwise statistics yield FC matrices that look different, do these matrices also have different topological and geometric features? Figure 2a shows the probability density of edge weights for each matrix (each column represents a pairwise interaction statistic, following the order in Fig. 1). Some densities are highly skewed while others are more evenly distributed, suggesting differences in topological organization, such as the presence or non-presence of hubs, respectively. Figure 2b shows the weighted degree of every brain region in each of the FC matrices (brain regions *×* pairwise statistics). While there exist some patterns that are common to most pairwise statistics (e.g., high weighted degrees in dorsal attention, ventral attention, visual and somatomotor networks), there is also considerable variability across pairwise statistics. For instance, some families of statistics tend to have more spatially distributed hubs, additionally emphasizing transmodal regions, such as *precision*-based pairwise statistics that detect prominent hubs in default and frontoparietal networks (Fig. 2b).

**Figure 2.**
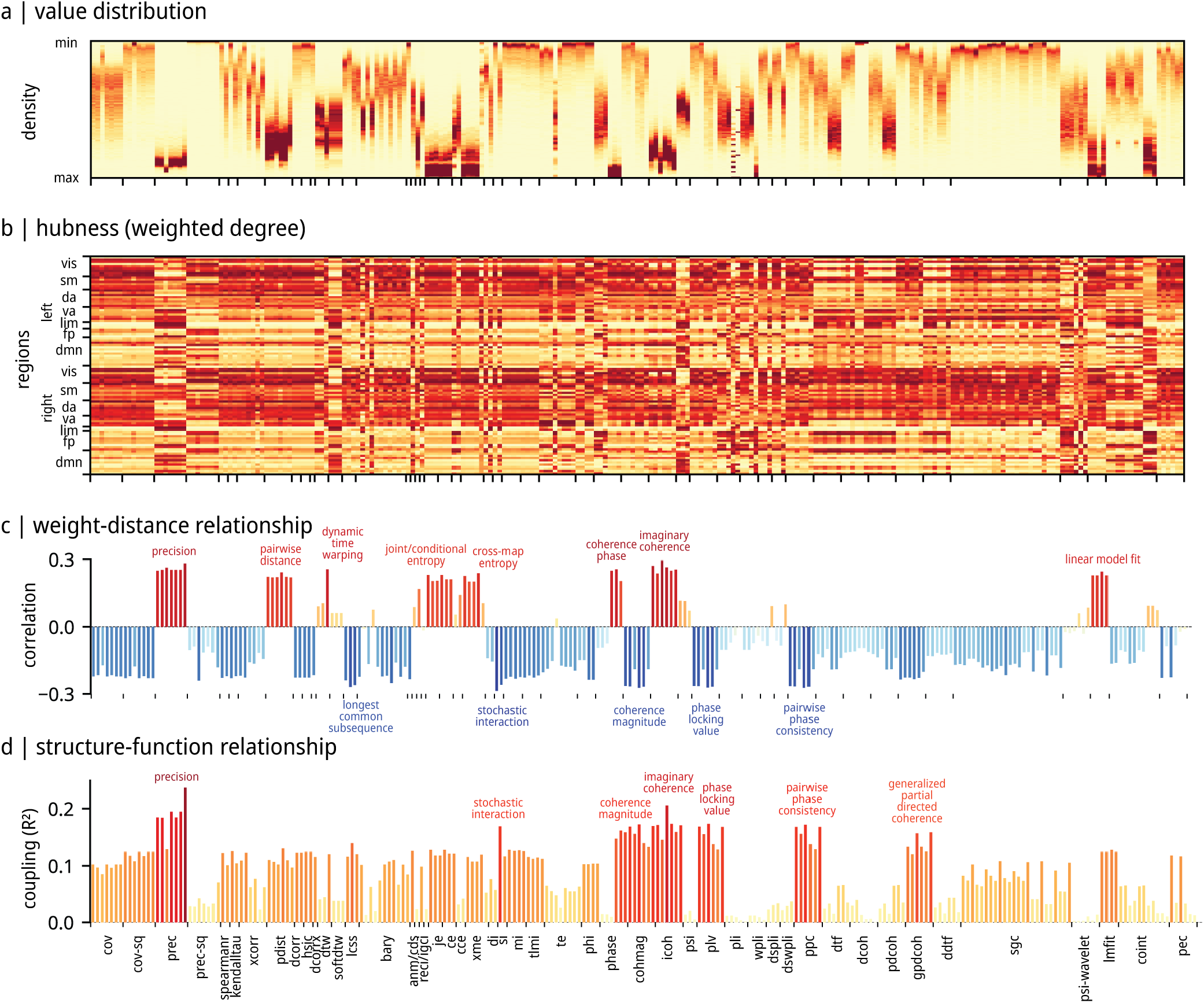
Benchmarking topological and geometric organization. **(a)** Value distribution for each interaction statistic. Values were min–max-normalized within each statistic. Darker red denotes greater density. **(b)** Ranking of hubs quantified by weighted degree (strength) of the pairwise statistic matrices. Absolute values are taken from the pairwise statistics before ranking. Note that pairwise statistics with positive correlations with spatial distance (shown in panel c) have flipped rankings to ensure a more consistent hub representation. Regions are ordered by intrinsic functional networks from [210] for left and right hemispheres. Darker red means greater weighted degree (“hubness”). VIS: visual, SM: somatomotor, DA: dorsal attention, VA: ventral attention, LIM: limbic, FP: fronto-parietal, DMN: default mode network. The organization of hubs when considering positive and negative values separately can be found in Fig. S2. The similarity of hub organization across pairwise statistics and their representation on the cortex are shown in Fig. S3. **(c)** Weight–distance relationship quantified by computing the Spearman’s rank correlation of each edge in each pairwise statistic matrix with inter-regional Euclidean distance (physical distance between brain regions). Colors and bar height represent the magnitude of correlation. Most extreme measures are labeled with text. **(d)** Structure–function coupling between matrices of interaction statistics and predictor matrices derived from structural connectivity. Structure–function coupling is represented using the coefficient of determination (adjusted *R*^2^), such that low values indicate poor structure–function coupling and high values indicate strong structure–function coupling [93, 199] (see *Methods* for details). Colors and bar height represent the magnitude of coupling. Most extreme measures are labeled with text.

Next, we quantify to what extent each of the pairwise statistics recapitulates two well-studied features of brain networks: (1) the inverse relationship between physical proximity and edge weight [19, 48, 74, 92, 181, 183], and (2) the positive relationship between structural connectivity and functional connectivity [72, 73, 92, 115, 183, 199]. Figure 2c shows, for each pairwise statistic, the correlation between the inter-regional Euclidean distance and the magnitude of functional connectivity. Note that some pairwise statistics are defined as the distance (dissimilarity) between time series (e.g., *precision, pairwise distance, linear model fit*); in those cases, greater values indicate dissimilar time series, and we expect to see a positive correlation between physical distance and functional connectivity. Overall, most pairwise statistics display a moderate inverse relationship between physical proximity and pairwise association (0.2 *<* |*r*| *<* 0.3), although several display a weaker relationship (|*r*| *<* 0.1). This finding illustrates how even a fundamental feature of brain networks that has been reported across imaging and tracing techniques, spatial scales, and species can vary substantially depending on how functional connectivity is defined. This suggests that pairwise statistics are differentially sensitive to different types of underlying mechanisms, a question we explore in greater detail in the *Decomposing FC matrices into information flow patterns* section.

Figure 2d shows, for each pairwise statistic matrix, the goodness of fit between diffusion MRI-estimated structural connectivity and the magnitude of functional connectivity. Here we expect a positive relationship, reflecting the fact that axonal projections support inter-regional signaling and the emergence of coherent dynamics among neuronal populations [183]. Again, we observe substantial variability across pairwise statistics, with structure–function coupling ranging from 0 *< R*^2^ *<* 0.25. Pairwise statistics with the greatest structure– function coupling include *precision, stochastic interaction*, and *imaginary coherence*. These results parallel the findings above in two ways. First, they show gross compatibility, but also substantial variability for an observation that has been reported in multiple studies. Second, we observe the strongest “expected” results (inverse relationship with distance, positive relationship with structural connectivity) for commonly used *covariance*-based pairwise statistics and for some others, like *precision*-based pairwise statistics. These statistics may be well-suited for optimizing structure–function coupling because they seek to partial out or account for shared influence among multiple regions, emphasizing functional interactions that arise from structural connections (see *Discussion*).

### Alignment with multimodal neurophysiological networks

The previous section demonstrates that even basic relationships with geometry and anatomical connectivity can vary substantially depending on how FC is estimated. Here we extend this question and consider how different types of FC correspond to other networks that reflect biological similarity between brain regions. Specifically, we estimate multiple forms of inter-regional similarity, including: correlated gene expression (Fig. 3a; derived from the Allen Human Brain Atlas microarray data), laminar similarity (Fig. 3b; derived from the Merker-stained BigBrain Atlas), neurotransmitter receptor similarity (Fig. 3c; derived from multiple PET tracers), electrophysiological connectivity (Fig. 3d; derived from magnetoencephalography), and metabolic connectivity (Fig. 3e; derived from dynamic FDG–PET). For a complete description of how each matrix is constructed, see *Methods*. Our main question here is how well each FC matrix aligns with inter-regional biological relationships estimated at different spatial and temporal scales.

**Figure 3.**
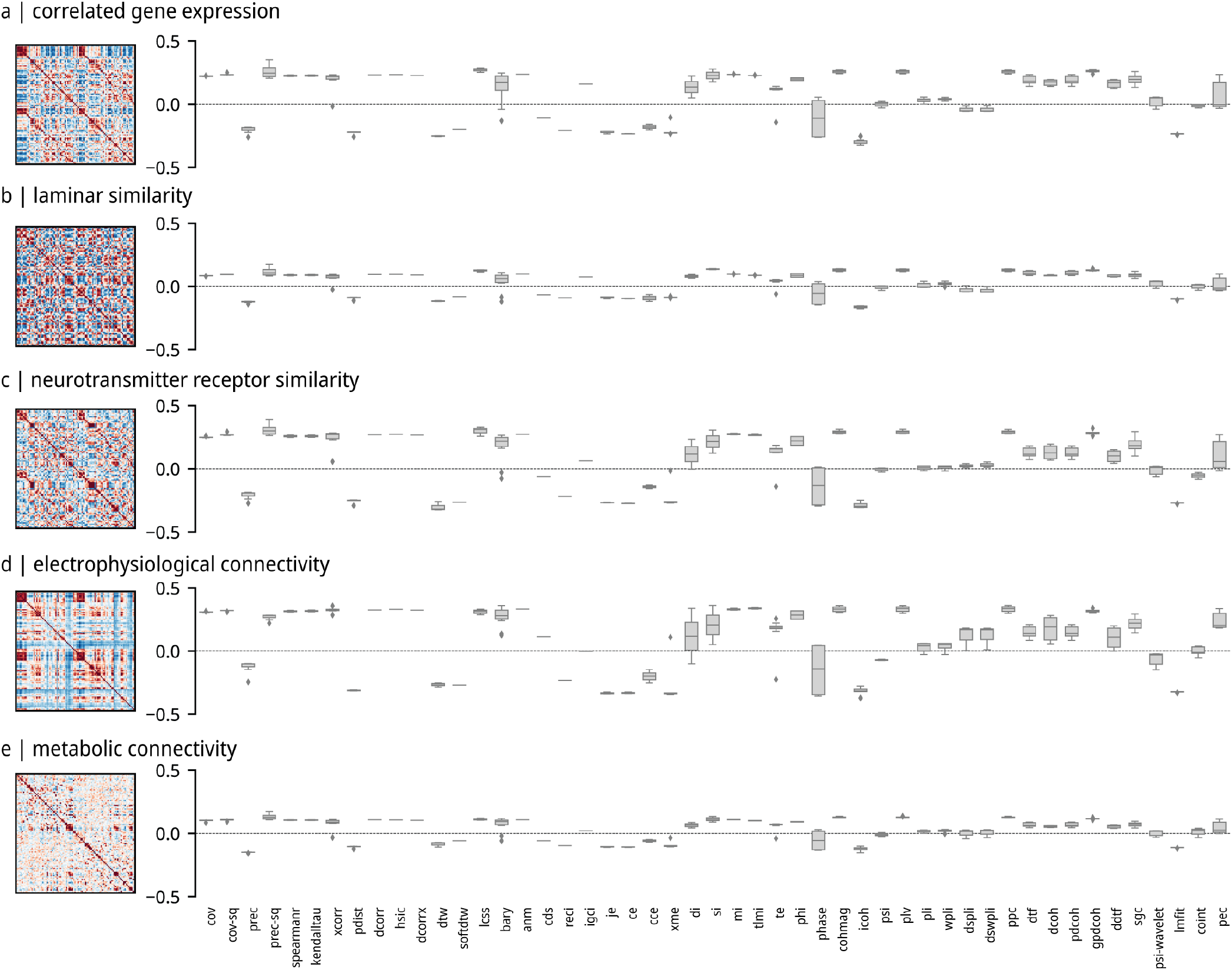
Alignment with multimodal neurophysiological networks. Inter-regional similarity networks (left) correlated with interaction statistics matrices (right) for **(a)** Correlated gene expression derived from Allen Human Brain Atlas microarray data [68, 105]. **(b)** Laminar similarity derived from BigBrain histological intensity profile segmented into cortical layers [4, 132, 201– 203]. **(c)** Neurotransmitter receptor similarity derived from PET tracer images of 18 neurotransmitter receptors and transporters [66, 106]. **(d)** Electrophysiological connectivity derived from resting-state MEG [67, 93, 167]. **(e)** Metabolic connectivity derived from correlating FDG-PET time-resolved activity [67, 79, 80, 200]. The y-axis represents the Spearman’s rank correlation between inter-regional similarity networks and interaction statistics matrices. The mean and variance of the alignment across the five neurophysiological networks are shown in Fig. S5.

Figure 3 shows the correlation between each FC matrix and each biological inter-regional similarity matrix. We observe the strongest correspondence with neurotransmitter receptor similarity and electrophysiological connectivity. This is consistent with the previous literature, and potentially reflects the fact that regions with similar chemoarchitectural profiles are subject to common neuromodulatory influences, leading to coherent electrophysiological dynamics [66, 93, 167]. We find similar results when we estimate the alignment between pairwise interaction statistics matrices and a “cognitive similarity” matrix that indexes how areas co-activate across cognitive tasks (derived from the Neurosynth meta-analytic engine) (Fig. S6). Perhaps counterintuitively, we do not observe strong correspondence between fMRI-estimated FC and FDG–PET-estimated metabolic connectivity, despite the fact that the two methods should theoretically be measuring related biological processes [200]. Finally, in what is a recurring theme, FC estimated using *precision*-based statistics generally continue to be closely aligned with multiple biological similarity networks.

### Quantifying individual differences

A common application of resting-state FC is to study individual differences [116]. Here we examine how FC estimated using different pairwise statistics can be used for (1) identifying individuals (“fingerprinting”) [3, 51], and (2) predicting individual differences in cognition and behavior [69, 127]. Figure 4a shows participant identifiability for FC matrices computed using different pairwise statistics. The identifiability index is a measure of effect size, so a magnitude of ≥ 0.8 is considered large [103]. Briefly, identifiability measures how similar an individual is to themselves across multiple scans, compared to other individuals [3, 103]. Consistent with previous reports, we find that *covariance* measures (e.g. Pearson’s correlation) generally perform well (identifiability ≈ 1.5) [103]. *Precision*-based statistics outperform all others (identifiability *>* 2.1), mirroring the results in the previous section. The broad question of whether FC organization persists across subjects and scans is sometimes altertatively formulated as test-retest reliability. For completeness, we additionally compute the intra-class correlation (ICC) using the four scans for each participant, and find similar results to fingerprinting (Fig. S7).

**Figure 4.**
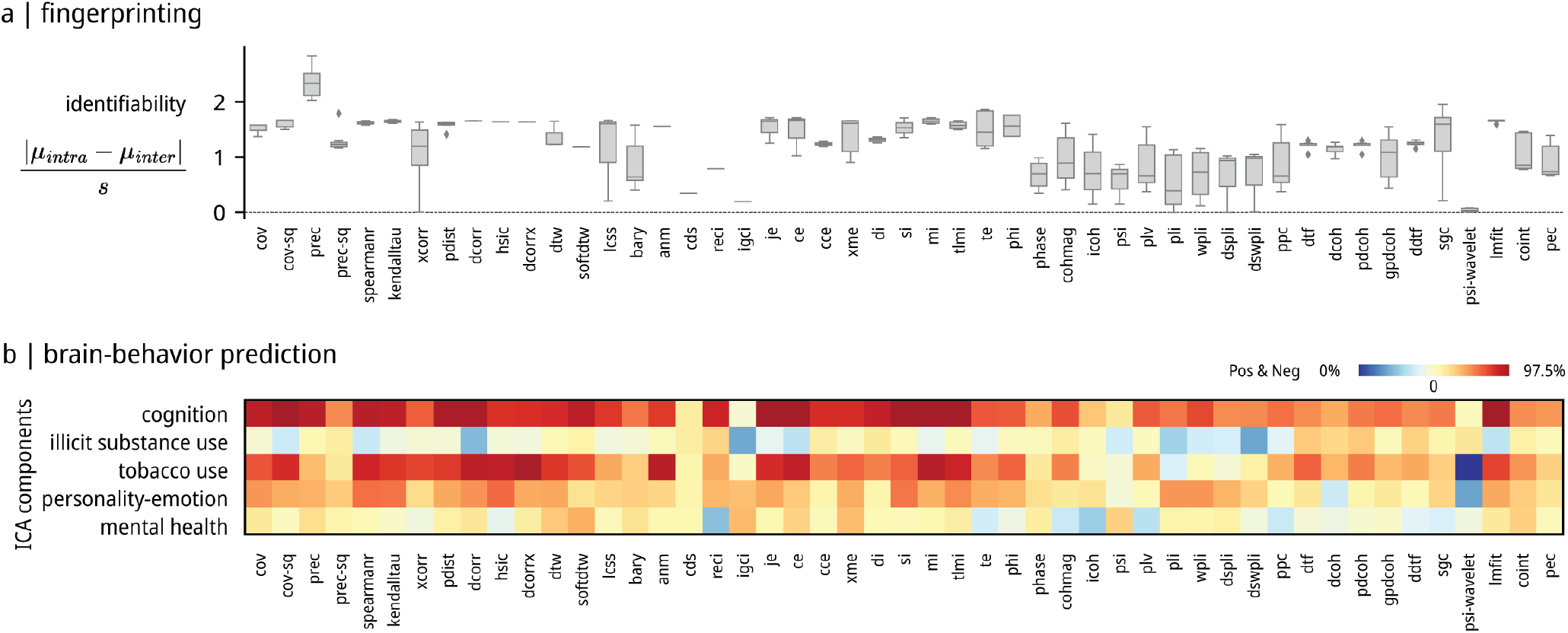
Quantifying individual differences. **(a)** Individual fingerprinting quantified using the identifiability index [3, 103], estimated from the 4 fMRI runs for each subject. **(b)** Brain–behavior prediction using pairwise statistics as predictors and 5 ICA-derived cognitive-behavioral components as the outcome [187]. Kernel ridge regression is performed under a nested 10-fold cross-validation setting. The heatmap colors display the mean Pearson’s correlation between the empirical and predicted behavior scores across the test folds. The colorbar covers both negative values (0th percentile to zero; in blue) and positive values (zero to 97.5th percentile; in red). Sensitivity analyses, using alternative machine learning algorithms (kernel ridge regression with a cosine kernel, linear ridge regression and LASSO regression) are shown in Fig. S8, Fig. S9, and Fig. S10.

We next consider how well different FC pairwise statistics can be used for out-of-sample prediction of individual differences in cognition and behavior. Following the approach outlined by Tian and colleagues [187], we apply independent component analysis to 109 measures in the HCP dataset to derive a five-component solution. The components broadly capture individual differences in cognition, illicit substance use, tobacco use, personality-emotion and mental health [187]. We then use kernel ridge regression in a nested 10-fold cross-validation setting to predict individual component scores from individual FC matrices [69, 86]. Figure 4b shows the mean correlation between empirical and predicted scores across the test folds. We generally observe greater prediction for cognition and tobacco use, and poor prediction for illicit substance use and mental health, consistent with previous reports [103, 165, 187]. Pairwise statistics that perform well for individual fingerprinting (e.g., *covariance, precision*, and information theory-based statistics) also tend to perform well for predicting cognition and behavior; likewise, pairwise statistics that perform poorly for fingerprinting also perform poorly here (e.g., spectral statistics). Collectively, the substantial variation in identifiability and prediction accuracy suggests that the choice of pairwise statistic for computing FC is an important one that could be tailored or optimized for different research questions.

### Decomposing FC matrices into information flow patterns

Up to now, we focused on associating FC matrices with other types of inter-regional relationships (e.g., structural connectivity, spatial proximity and inter-regional bi-ological similarity), and with exogenous measures (e.g., individual identity or behavior). Here we ask whether FC computed using different pairwise statistics reflects different underlying patterns of information flow. We estimate, for instance, “synergistic” interactions where two sources of information, when considered together, provide new information that cannot be retrieved from either source individually, and in contrast, “redundant” interactions where the opposite is true, and each source provides the same information as the other. A recent information-theoretic framework makes it possible to partition pairwise interactions into synergistic, redun-dant, and unique information, also known as “information atoms” [62, 100, 101, 113, 195, 205] (Fig. 5a).

**Figure 5.**
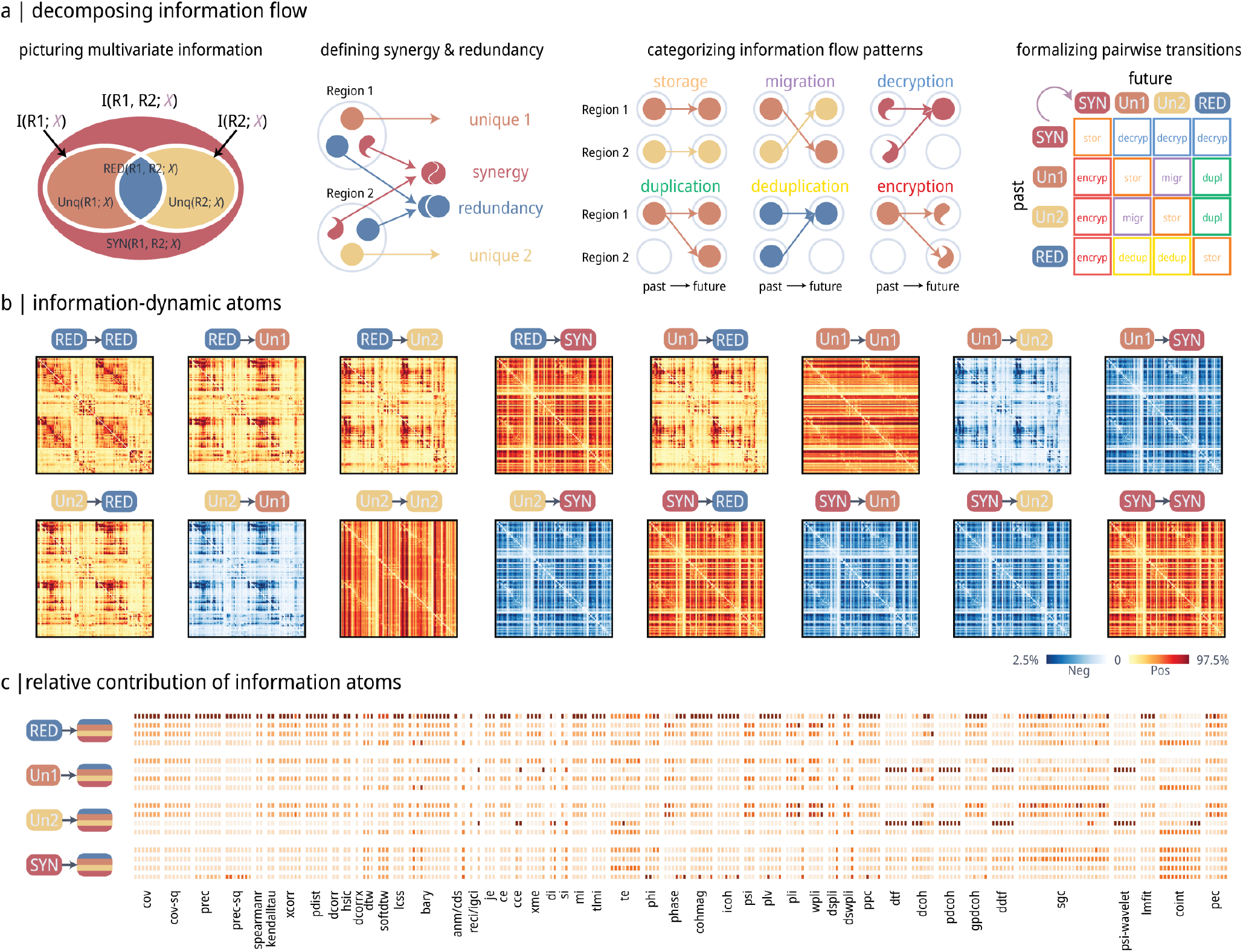
Decomposing FC matrices into information flow patterns. **(a)** Schematic of the Integrated Information Decomposition framework (ΦID), showing how pairwise information is estimated and decomposed into basic motifs of information flow [100, 101, 113]. **(b)** The framework yields 16 distinct information-dynamic atoms that represent different patterns of information flow. **(c)** Relative contribution of the 16 information-dynamic atoms to each pairwise interaction statistic, quantified as relative importance estimated using dominance analysis [30] (see *Methods*). In each column, darker red shows greater contribution from a specific information-dynamic atom. The 16 rows, grouped by the “past” state, correspond to the 16 information-dynamic atoms in the same order as panel b.

Briefly, for each pair of cortical regions (treated as sources) we can ask how much information about their future BOLD activity can be obtained from knowing their past activity—and whether this information is carried redundantly by each of them separately, or uniquely by one of them, or synergistically by both together. We can then also ask if the way that information is carried changes over time, giving rise to different types of information dynamics. For example, if information was initially provided uniquely by region A, and then it is pro-vided uniquely by region B, this is a case of *information transfer* from A to B.

Figure 5b shows the 16 information flow patterns arising from this information decomposition ([113]; see *Methods* for details). We then estimate the contribution of each of the 16 information flow patterns to each FC matrix (Fig. 5c). We find that classic statistics, such as *covariance, precision* and *mutual information*, mostly reflect the pattern whereby redundant information stays as redundant. Some spectral statistics, such as *directed transfer function* and *partial coherence*, predominantly reflect a pattern where information that is exclusively provided by one region stays as unique to that region. While both of the cases above belong to a pattern of information storage, whereby information is consistently conveyed in the same way over time, a greater diversity of information flow patterns exists. For example, we observe the presence of information migration, duplication, and deduplication in *phase lag value*. We also observe information encryption and decryption (also known as downward and upward causation, see [113]) in *transfer entropy* and *cointegration*. Altogether, these results show that, while most statistics capture redundant information storage, there exists a wider landscape of information flow patterns that can potentially be selectively sampled using specific pairwise statistics.

### Sensitivity analyses

We finally seek to determine the extent to which these results are sensitive to the several processing and data-handling choices that exist in resting state fMRI network modelling. We first test the stability of the group-level similarity matrix (originally shown in Fig. 1). We perform 1000 random splits of the sample into *Discovery* and *Replication* sets and compute the correlation between them. Figure 6 shows that the distribution of correlation coefficients is centered above *r* = 0.999. To test the effect of atlas, we compute the similarity between matrices generated using a functional parcellation (Schaefer 100*×*7; [161]) and an anatomical parcellation (Desikan–Killiany; [41]), revealing a correlation of *r* = 0.96. To test the effect of atlas resolution, we compute the similarity between matrices generated using a lower-resolution atlas (Schaefer 100*×*7) and a higher-resolution atlas (Schaefer 200*×*7), revealing a correlation of *r* = 0.98. Finally, we test the effect of global signal regression by computing the similarity between matrices generated with and without global signal regression, revealing a correlation of *r* = 0.82. Altogether, these sensitivity checks suggest that the global relationships among pairwise statistics are relatively stable with respect to multiple methodological choices.

**Figure 6.**
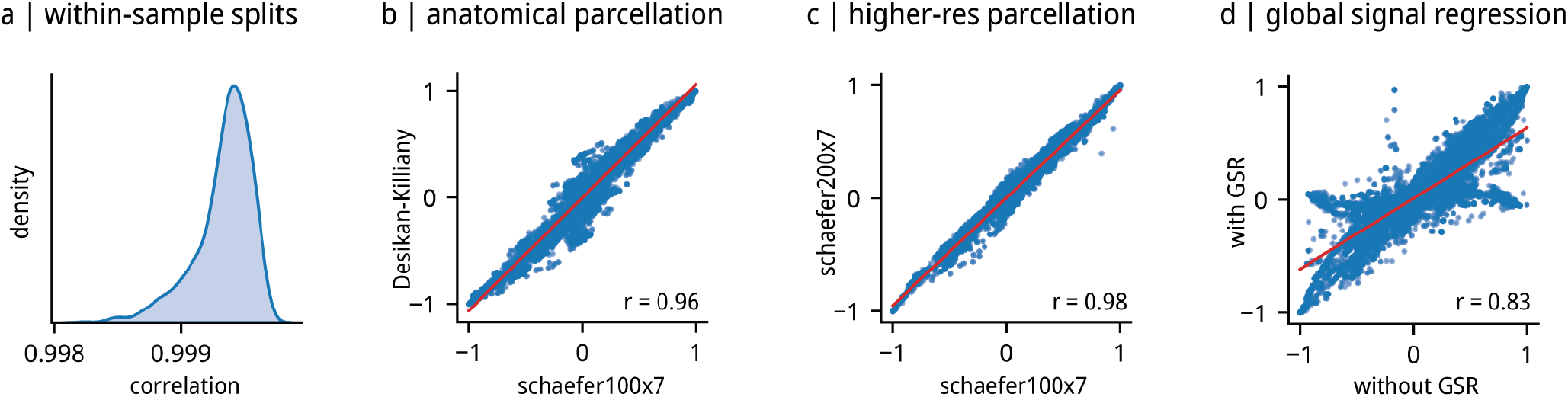
Sensitivity analyses. The group average similarity profile of the pairwise statistics in Fig. 1 is reproduced under different processing conditions for comparison. **(a)** Distribution of correlations between split-half discovery and validation sets for 1000 random splits. **(b)** Correlation between the functionally derived atlas (Schaefer 100-node 7-network; [161]) used in the main analyses and an anatomically derived atlas (Desikan–Killiany; [41]). **(c)** Correlation between the 100-node atlas (Schaefer 100-node 7-network, Schaefer100*×*7) used in the main analyses and a higher-resolution atlas (Schaefer 200-node 7-network, Schaefer200*×*7). A subset of 179 interaction statistics were used in this panel for faster calculation, and can be found in Table S2. **(d)** Correlation between functional time series without global signal removal used in the main analyses and with global signal removal. GSR: global signal regression. Additional analyses regarding the effect of participant motion can be found in Fig. S11). Spearman’s rank correlation is used in all panels.

## DISCUSSION

Resting state FC is rapidly becoming one of the most widely used brain imaging phenotypes. Despite its popularity, the operational definition of FC is arbitrary, and most groups use simple zero-lag linear correlations by default. In the present report we benchmark the network architecture, biological underpinnings, and brain– behavior associations of FC matrices computed using a large library of pairwise interaction statistics. Our results reveal a rich landscape of methods that are sensitive to different features of brain organization.

Even for well-studied phenomena, we observe substantial variability across methods. The arrangement of highly connected hub regions, a topic of great interest over the past 10–15 years [139, 179, 192], systematically varies depending on method, with some localizing hubs in unimodal cortex and others more widespread across the unimodal–transmodal axis. The weight–distance relationship, reported not only for FC-fMRI but also in diffusion MRI [20, 145], and tract-tracing in multiple species [184, 190], is captured by most methods, but the magnitude of the effect varies considerably. Finally, a similar result is observed for structure–function coupling, whereby most methods identify an overall positive relationship, but the effect size displays variability across methods. In other words, the choice of pairwise interaction statistics has substantial influence on the spatial and topological organization of reconstructed functional networks.

One reason for the observed variability is that pairwise statistics are sensitive to different underlying mechanisms of inter-regional signaling [169]. We find that different FC methods often align with different forms of inter-regional biological similarity, from microscale correlated gene expression or receptor similarity, to macro-scale electrophysiological coupling. Indeed, numerous reports have found evidence of association between resting-state BOLD FC and correlated gene expression [144], receptor similarity [66] and electrophysiological rhythms [93, 104, 167]. Indeed, the different pairwise statistics are optimized to capture different types of communication processes [63, 85, 180]. Resting-state functional dynamics are thought to be mostly macroscopically linear [126], and as a result, many conventional FC methods are designed to capture linear effects. However, the complexity of functional dynamics extends beyond simple linear effects and a broader set of pairwise statistics is necessary to completely capture the rich spectrum of interactions in fMRI BOLD neural dynamics [26, 29, 32].

Ultimately, one of the main reasons why neuroscientists study statistical relationships between regional BOLD time-series is the belief that brain regions exchange, store, and process information, and that this information can be reflected by statistical relationships. However, there is growing understanding that information can be transmitted, processed, and stored in different ways—raising the question of how each pairwise statistic captures (or fails to capture) these different kinds of information dynamics. To directly address this question, we applied information decomposition and found that different FC methods align with different forms of information dynamics. Most FC methods appear to capture storage of redundant information, whereby both regions convey the same information—as previously observed for Pearson correlation [100, 196, 197]. However, some measures are sensitive to other forms of communication, including synergistic and unique information flow. These results demonstrate a multitude of communication patterns between brain regions that are explored less often, but which should be taken into account for a more comprehensive mapping of the functional connectome and more nuanced inferences about what functional connectivity represents [98, 129, 133,150].

To summarize our benchmarking findings, Table S5 shows the ranking of pairwise statistics according to six (potentially desirable) criteria: (1) negative weight– distance relationship, (2) positive structure–function coupling, (3) close correspondence with biological similarity networks, (4) high individual–participant identifiability, (5) high brain–behavior prediction and (6) low susceptibility to participant motion. Collectively, these results suggest that there is not necessarily a single optimal pairwise statistic, but rather different options that can be used to target desired mechanisms. In this sense, our results can be seen as a rough guide for matching a pairwise statistic to an experimental question. A salient example is how the choice of FC method is context-dependent in individual differences and brain-behavior relationships, where we find that the predictive utility of an FC method depends on the phenotype that one seeks to predict [103, 165, 187]. More broadly, our results highlight the idea that, in the absence of any ground truth, picking a pairwise statistic is an important ques-tion that strongly depends on the research question at hand [71, 97, 98, 129, 133, 143, 150].

What recommendations can be derived from the present findings? Although we sampled a limited set of possible analyses, some broad arcs come into focus. First, as discussed above, a pairwise statistic should be matched to the experimental question. Second, covariance (distance)-based methods appear to have many desirable properties, including robust relationships with physical proximity, structural connectivity, and biological inter-regional similarity, as well as the capacity to differentiate individuals and predict individual differences in multiple phenotypes. Methods based on *precision* (inverse covariance or partial correlation) stand out. Indeed, these measures have often been touted as the superior alternative to the Pearson’s correlation for estimating FC [47, 64, 107, 108, 110, 185]. By removing mutual dependencies on common influences from other areas, precision has the theoretical advantage of more directly measuring directed, anatomically-mediated interactions among brain areas [38, 50, 77, 94, 95, 135, 143, 152, 162, 176, 198]. An exciting future avenue would be to combine multiple FC matrices to engineer new types of FC that are potentially sensitive to a wider range of desirable properties [67].

Finally, the present results should be interpreted in light of multiple methodological limitations. First, we only considered undirected components of pairwise statistics, effectively ignoring directed or causal mechanisms [143, 208]. Second, although we ensured robustness to common preprocessing choices such as parcellation type and size, and removal of the global signal, we did not exhaustively consider the effects of all processing choices, including acquisition parameters, and alternative motion and artifact correction methods [102]. Third, we did not exhaustively consider all common research questions, such as the lifespan trajectory of FC or the effects of psychiatric and neurological disease on FC [147]. Fourth, we only focused on descriptive pairwise interaction statistics, and did not explicitly consider model-based “effective connectivity” methods, such as structural equation modeling or path analysis, dynamic causal modeling or biophysical neural mass modeling [53, 54, 56, 96, 110, 112, 168, 206].

In summary, the present report comprehensively benchmarks the architecture of resting state BOLD FC using a large library of pairwise statistics. We observe substantial variation across FC methods and across a wide array of analyses, reflecting differential sensitivity to biological features and to types of information flow. As FC continues to grow in popularity as a neuroimaging phenotype, our results provide the foundation for future studies to tailor their choice of FC method to the neurophysiological mechanism they are targeting and to their research question.

## METHODS

### Resting-state functional MRI

Resting-state functional time series from 326 unrelated subjects were obtained from the Human Connectome Project Young Adults cohort (HCP-YA; S1200 release [194]). Structural and functional MRI data were preprocessed using HCP minimal preprocessing pipelines [59, 193]. High-resolution T1w and T2w structural images were corrected for gradient distortion and registered to MNI152 atlas. Cortical surfaces were constructed using FreeSurfer recon-all procedure. Resting-state BOLD functional images (4 scans of approximately 15 minutes long for each subject) were corrected for slice timing, gradient distortion, motion, EPI distortion, and registered to the high-resolution T1w structural image. It further underwent intensity normalization and bias removal. The surface representations were then created by mapping the volumetric BOLD signal to the fsLR gray-ordinate space using MSMAll, a multi-modal surface-based functional alignment algorithm [146]. Physiological noise and confounds were removed with the ICA-FIX procedure [155]. Details of the preprocessing steps can be found in the original technical reports [59].

### Calculating pairwise interactions with *pyspi*

We used the recently developed Python Toolkit of Statistics for Pairwise Interactions (*pyspi*; v0.4.1, commit c19d06) to calculate the alternative measures (statistics of pairwise interactions; SPIs) of functional connectivity [32]. Resting-state fMRI time series derived in the previous step were parcellated using Schaefer 100-node 7-network atlas [161] and normalized (z-scored along the time dimension) before *pyspi* calculation. Starting with the original list of SPIs, we derived a subset of SPIs with reasonable calculation time (< 30 minutes) for a single subject and calculated the SPIs for all individual subjects and resting-state runs. After aggregating the results, we further excluded the SPIs with 1) zero variance, or 2) infinity or NaN values for at least 1/4 of all subjects and runs, finally getting 239 SPIs from 49 pairwise interaction measures across 6 major categories (see Table S1 for the full list of SPIs used).

The calculation resulted in 239 node-by-node matrices for each subject and run. A group consensus matrix was calculated for each statistic by taking the average across all subjects and runs (shown in Fig. 1; see Fig. S1 for variance). A total of 239 group consensus matrices were generated, which we refer to them as group-averaged measure matrices. We also calculated the similarity of the statistics by taking the Spearman’s rank correlation between pairs of statistic matrices for each subject and run, which we refer to as similarity profile matrices. A group consensus similarity profile matrix was calculated by taking the average across subjects and runs.

Unless otherwise noted, we used the upper triangu-lar values for the analyses (see Fig. S12 for a brief account of directed pairwise statistics), and using Spearman’s rank correlation coefficient to assess the relationships between SPIs and other measures. The calculation was performed using a singularity container available (https://osf.io/75je2/) for reproducibility.

### Structure–function relationship

#### Structural network reconstruction

Structural network of the cohort was reconstructed from diffusion MRI tractography. Diffusion MRI scans were processed using MRtrix3 package [189]). Fiber orientation distributions were modelled using multi-shell multi-tissue constrained spherical deconvolution algorithm [42, 81]. White matter streamlines were then reconstructed [188] and optimized [173] to provide robust estimate of tract weights. We estimated a binary group consensus structural connectivity matrix using a distant-dependent algorithm, that approximates the group-level average edge length distribution [21]. The final weighted group consensus matrix was then calculated by applying the binary matrix on the simple average of structural connectivity matrices of all subjects.

#### Structure–function coupling estimation

Following previous practices of quantifying structure– function relationship [93, 199], we used a multilinear regression model with network communication predictors to quantify the correspondence between structural and functional networks. This approach takes into account of potential dynamics processes happening on the network and provides a multi-faceted view of structure– function correspondence than using the structural connectivity alone. We adopted Euclidean distance and five commonly used network communication measures derived from the group consensus structural connectivity matrix as predictors. They represent a spectrum of routing strategies ranging from centralized, globally optimized shortest path to decentralized, locally focused diffusion [6–8, 164]. We estimated the goodness of fit, adjusted *R*^2^ to quantify the extent of structure–function coupling in this case.

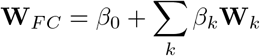

where **W**_*F C*_ denotes the pairwise interaction measures and **W**_*k*_ denotes predictor matrices: Euclidean distance, shortest path length, navigation efficiency, search information, communicability, and diffusion efficiency.

We additionally calculated a more direct version of structure–function coupling using Spearman’s rank correlation between non-zero elements of the structural connectivity and the pairwise interaction statistic matrices (Fig. S4) [14].

#### Network communication measures

We used *Euclidean distance* between region centroids as the physical distance between nodes. We also derived a *connection length matrix L* from the structural connectivity matrix when it comes to quantify the cost of traversing the edges. We used a monotonic weight-to-length transform in the form of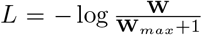. The resulted connection length matrix (*L*) will have infinity values between a pair of regions that do not have a direct structural connection.

*Shortest path length* represents the shortest distance to travel from a source and a target node [82]. We calculated the shortest path lengths using Floyd–Warshall algorithm [52] with connection length matrix *L*.

*Network navigation* was introduced to brain networks by Seguin and colleges [25, 119, 163, 166], quantifying routing without global optimization by simulating a walker that steps towards the neighbor node that is closest in distance to the target node. Here we used Euclidean distance as the distance metric and calculated navigation efficiency is calculated as the inverse of navigation path length.

*Search information* measures the amount of information necessary for a random walker on the network to travel along a specific path and does not take detours. The measures were adapted to the brain networks and the shortest path on weighted network in [60, 61, 149, 163].

*Communicability* measures the amount of possible routes between a source and a target node pair. It is defined as the weighted sum of all paths and walks between those nodes [37, 49].

*Diffusion efficiency* is calculated as the inverse of the mean first passage time, which quantifies the time (number of steps) expected for a random walker to travel from a source to a target node. For asymmetric measures, we symmetrized the matrix by taking the average of the matrix with its transpose [60, 61, 123].

The network measures were implemented using the *Brain Connectivity Toolbox* ([151]; https://sites.google.com/site/bctnet, Version 2019-03-03), *Brainconn* (https://github.com/FIU-Neuro/brainconn, master branch at commit 8cd436), and *netneurotools* (https://github.com/netneurolab/netneurotools, v0.2.3).

### Biological networks

We adopted annotated networks from multiple modalities to contextualize the functional relationships. Here, we briefly describe how we acquire the networks. More technical details can be found in the previous reports [67].

*Electrophysiology connectivity* was derived from resting-state magnetoencephalography (MEG) recordings [10]. Resting-state MEG data (approximately 6 minutes for each subject) for *N* = 33 healthy unrelated subjects were taken from Human Connectome Project (HCP). Preprocessing was carried out using open-source Brainstorm software (https://neuroimage.usc.edu/brainstorm/; [186]). Briefly, raw MEG recordings were registered to high-resolution anatomical space before submitted to notch filtering (60, 120, 180, 240, 300 Hz), high-pass filtering (0.3 Hz), band channel removal, automatic artifact removal. Artifacts including heartbeats from electro-cardiogram (ECG), eye blinks from electrooculogram (EOG), saccades, muscle movements as low-frequency (1 to 7 Hz) and high-frequency (40 to 240 Hz) components, and noisy segments) were removed using Signal-Space Projections (SSPs). Sensor-level data was then submitted to source estimation using linearly constrained minimum variance (LCMV) beamformer on HCP fsLR4K surface. The “median eigenvalue” method from Brainstorm was used to reduce the variable source depth effect. Time series on fsLR4k surface were parcellated to Schaefer100*×*7 atlas using the first principal component of the corresponding vertices. MEG functional connectivity matrices were estimated using amplitude envelope correlation [28] for the 6 canonical frequency bands: delta (*δ*; 2 to 4 Hz), theta (*θ*; 5 to 7 Hz), alpha (*α*; 8 to 12 Hz), beta (*β*; 15 to 29 Hz), low gamma (lo-*γ*; 30 to 59 Hz), and high gamma (hi-*γ*; 60 to 90 Hz). Spatial leakage effect was corrected using an orthogonalization process [34]. The final electrophysiology connectivity matrix used in this project is derived as the first principal component of the connectivity matrices for the 6 canonical bands. Details of preprocessing can be found in [67, 167].

*Correlated gene expression network* quantifies the transcriptional similarity between cortical regions. Spatially resolved microarray gene expression data were obtained from the Allen Human Brain Atlas (AHBA) [68], preprocessed and mapped to the Schaefer100*×*7 atlas using the abagen toolbox [105]. Briefly, the preprocessing procedure includes intensity-based filtering, representative probe selection, tissue sample matching, normalization, and aggregation [5]. The final region-by-region correlated gene expression matrix was estimated by calculating the Pearson’s correlation coefficient using normalized gene expression profiles across regions.

*Laminar similarity network* measures the similarity of cellular profiles across the cortical layers between pairs of regions. Histology-based cell-staining intensity values were derived from a postmortem brain, quantifying cell density and soma size [201–203]. Depth-resolved intensity values were sampled from 50 equivolumetric surfaces from white to pial surface. The intensity profiles were acquired on *fsaverage* surface using BigBrain-Warp toolbox [4, 132], and subsequently parcellated to Schaefer100*×*7 atlas. The region-by-region laminar sim-ilarity network was calculated using partial correlation, correcting for mean intensity across cortical regions.

*Metabolic connectivity* represents the co-fluctuation of glucose metabolism between cortical regions. Volumetric FDG-PET ([^18^F]-fluorodeoxyglucose) images were recorded over time for 26 healthy subjects [79, 80]. PET images were reconstructed and preprocessed using a previously reported pipeline, resulting 225 16s fPET volumes for each recording [200]. They are subsequently motion corrected, underwent a spatial temporal gradient filter, and registered to the MNI152 template. Finally, they are parcellated to Schaefer100*×*7 atlas and metabolic connectivity matrix was calculated as Pearson’s correlation coefficient for each subject. The group-averaged matrix was used in this project.

*Receptor similarity network* measures the similarity of receptor density profiles between regions. PET tracer data for 18 neurotransmitter receptors and transporters were taken from [66] and *neuromaps* (v0.0.1, https://github.com/netneurolab/neuromaps [106]). The neurotransmitter systems include dopamine (D_1_ [83], D_2_ [158, 170, 172, 211], DAT [45]), norepinephrine (NET [17, 31, 43, 156]), serotonin (5-HT_1A_ [160], 5-HT_1B_ [11, 57, 109, 117, 118, 136, 159, 160], 5-HT_2_ [18], 5-HT_4_ [18], 5-HT_6_ [140, 141], 5-HTT [18]), acetylcholine (*α*_4_*β*_2_ [11, 70], M_1_ [120], VAChT [2, 16]), glutamate (mGluR_5_ [44, 171]), GABA (GABA_A_ [124]), histamine (H_3_ [58]), cannabinoid (CB_1_ [46, 121, 125, 142]), and opioid (MOR [84]). Each PET image was parcellated to Schaefer100*×*7 atlas. The final receptor similarity matrix was calculated as Pearson’s correlation coefficient between the receptor profiles for pairs of regions.

### Fingerprinting

Fingerprinting of individual differences were calculated using the identifiability metric proposed in [3, 103].

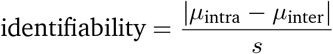

For each pairwise statistic, 4 matrices from BOLD runs per subject were used to calculate the mean values of within-subject correlations *µ*_intra_ and between-subject correlations *µ*_inter_. The pooled standard deviation *s* is also estimated. The resulting measure of identifiability is analogous to an effect size statistic [103].

### Behavior prediction

We used a robust set of ICA-derived cognitive-behavioral phenotypes derived by [187]. Briefly, HCP behavioral dataset were filtered for measures related to alertness, cognition, emotion, sensory–motor function, personality, psychiatric symptoms, substance use, and life function. A total of 109 measures were selected and subjected to an independent component analysis (ICA) procedure. Before the ICA procedure, normalization (87 out of 109) and confound regression (age, sex) was carried out to clean the raw behavioral data. Consistency and reliability of the ICA procedure was validated with bootstrapping, agglomerative clustering, followed by a sampling and matching process. A five-component model emerged as the most robust and concise representation of the original data structure: cognitive performance, illicit substance use, tobacco use, personality and emotion traits, and mental health. Details of the can be found in [187]. The intersecting *N* = 310 subjects were used for this study.

For the pairwise interaction measures, we used the vectorized upper triangular values of the SPI matrices for each subject, averaged across the 4 BOLD runs. To make the prediction more robust, we filtered the data using Quartile Coefficient of Dispersion (QCoD) to provide a conservative representation of the predictor vector. We first calculated QCoD across subjects for each SPIs and excluded those with minimal variance for all region pairs (Absolute max QCoD < 0.01; *pli_multitaper_max_fs-1_fmin-0_fmax-0-25, pli_multitaper_max_fs-1_fmin-0-25_fmax-0-5, wpli_multitaper_max_fs-1_fmin-0_fmax-0-25*). For each prediction, we further calculated the 10 and 90 QCoD percentile and included only the region pairs within this range to avoid spurious values with very large or little variance that may affect the prediction.

Following previous best practices [69, 86, 90, 127], we used kernel ridge regression with linear kernel for behavior prediction. We set up the prediction pipeline with nested k-fold cross validation. The inner 10-fold cross validation loop was used to select the optimal regularization parameter *α*, and the final performance was evaluated in the independent test split in the outer 10-fold cross validation loop. Both training and testing data were standardized using statistics estimated only from the training data to avoid leakage. We calculated Pearson’s correlation between empirical and predicted values for the final evaluation. The same process was also repeated with kernel ridge regression with cosine kernel, linear ridge regression, and LASSO regression. A comparison of average performance and variability across the 49 pairwise measures are show in Fig. S8 and Fig. S9, respectively. The performance details for each of the 239 individual statistics are shown in Fig. S10.

### Integrated information decomposition (ΦID)

We used Integrated Information Decomposition (ΦID; [100, 101, 113]) a temporally extended framework of Partial Information Decomposition (PID; [62, 78, 205]) to estimate the information flow patterns (“information flow atoms”).

The original PID framework aims to study the multivariate information by jointly considering multiple source variables with an additional target variable. As shown in Fig. 5a, in a 2-variable scenario, *I*(*R*_1_; *X*), *I*(*R*_2_; *X*), and *I*(*R*_1_, *R*_2_; *X*) represent their specific information, quantifying the information provided by the source variables when provided the information about the target variable *X*. PID decomposes the information contents into their unique information component (Unq(*R*_1_; *X*), Unq(*R*_2_; *X*)), a redundant information component (RED(*R*_1_, *R*_2_; *X*)), and a synergistic information component (SYN(*R*_1_, *R*_2_; *X*)). Here, the term “redundant” suggests information identically provided by each of the two variables individually, and the term “synergistic” suggests new information that emerges when the two variables are considered together (see [78, 101, 113, 205] for formal definitions).

ΦID extends this framework by introducing a temporal dimension. Taking a pair of time series as inputs, ΦID defines a past and a future state, and derives 16 information flow atoms, denoted as pairwise transitions between the initial 4 information atoms. The 16 types of information flow can be mechanistically categorized into several types: storage (information that remains carried in the same way over time; `Red → Red, Un`^1^ → `Un`^1^, `Un`^2^ → `Un`^2^, and `Syn → Syn`), duplication (information that becomes redundantly available from both variables, and was not before; Un^1^ → Red, and Un^2^ → Red), migration (information that moves between variables, such that it was uniquely present in a single variable, and subsequently it is uniquely present in the other; Un^1^ → Un^2^ and Un^2^ → Un^1^), deduplication (information that is pruned from duplication, such that it is no longer redundant; Red → Un^1^ and Red → Un^2^), decryption (collective/distributed information that becomes individual information in the future, also known as “downward causation”; Syn → Un^1^, Syn → Un^2^, and `Syn → Red`), and encryption (individual information that becomes collective/distributed information in the future, also known as “upward causation”; Un^1^ → Syn, Un^2^ → Syn, and `Red → Syn`) (see [100, 101, 113] for more rigorous definitions).

Technically, ΦID requires a choice of how redundancy is defined, just like PID. Here, we chose the minimum mutual information (MMI) definition of redundancy, following previous work [12, 99, 100, 113]. Overlapping segments of the functional time series with one time step (TR) delay were used to define the past and future states. We calculated ΦID for every pair of the original functional time series using time-delayed mutual information (mutual information between the past and future states) under the Gaussian assumption for continuous variables. This process generated 16 information flow matrices (Fig. 5b). Note that there are many potential implementations of redundancy and temporal states; here we adopt a straightforward definition as previously validated in [100, 101, 113]. An open-source implementation can be found at https://github.com/Imperial-MIND-lab/integrated-info-decomp.

In order to establish the relationship between information flow atoms and pairwise interaction statistics (Fig. 5c), we constructed linear models utilizing the former as predictors and the latter as the outcome. We used Dominance Analysis [9, 30] to quantify the contribution of individual predictors in the presence of potential multicollinearity [88]. The “total dominance” statistic is used to calculate the relative contribution of each predictor compared to the goodness of fit (*R*^2^) of the full linear model. The function is implemented in *netneurotools* (https://github.com/netneurolab/netneurotools), which is adapted from the *Dominance-Analysis* package (https://github.com/dominance-analysis/dominance-analysis).

### Sensitivity analyses

For sensitivity analyses, the time series were additionally parcellated into Desikan-Killiany atlas [41] and Schaefer 200-node 7-network atlas [161]. They also underwent global signal removal. To effectively calculate the sensitivity analysis using a higher atlas resolution with 200 regions, we generated a minimized list of SPIs by removing those taking more than 30 minutes to calculate for a single subject, resulting in 197 SPIs calculated. After taking the intersection with the list of SPIs above, 179 SPIs were used for the sensitivity analysis (see Table S2 for the full list of SPIs used). The group-average measure similarity matrix shown in Fig. 1 was calculated for each scenario and Spearman’s rank correlation coefficient was used to quantify the correlation.

## ACKNOWLEDGMENTS

We thank Vincent Bazinet, Filip Milisav, Eric Ceballos and Asa Farahani for comments and suggestions on the manuscript. BM acknowledges support from the Natural Sciences and Engineering Research Council of Canada (NSERC), Canadian Institutes of Health Research (CIHR), Brain Canada Foundation Future Leaders Fund, the Canada Research Chairs Program, the Michael J. Fox Foundation, and the Healthy Brains for Healthy Lives initiative. ZQL acknowledges support from the Fonds de Recherche du Québec – Nature et Technologies (FRQNT). AIL acknowledges the support of the Natural Sciences and Engineering Research Council of Canada (NSERC), [funding reference number 202209BPF-489453-401636, Banting Postdoctoral Fellowship] and FRQNT Strategic Clusters Program (2020-RS4-265502 - Centre UNIQUE - Union Neuroscience & Artificial Intelligence - Quebec) via the UNIQUE Neuro-AI Excellence Award. BTTY is supported by the NUS Yong Loo Lin School of Medicine (NUHSRO/2020/124/TMR/LOA), the Singapore National Medical Research Council (NMRC) LCG (OFLCG19May-0035), NMRC CTG-IIT (CTGIIT23jan-0001), NMRC STaR (STaR20nov-0003), Singapore Ministry of Health (MOH) Centre Grant (CG21APR1009), the Temasek Foundation (TF2223-IMH-01), and the United States National Institutes of Health (R01MH120080 & R01MH133334). Any opinions, findings, and conclusions or recommendations expressed in this material are those of the authors and do not reflect the views of the funders.

## Supplemental Materials

**Figure S1.**
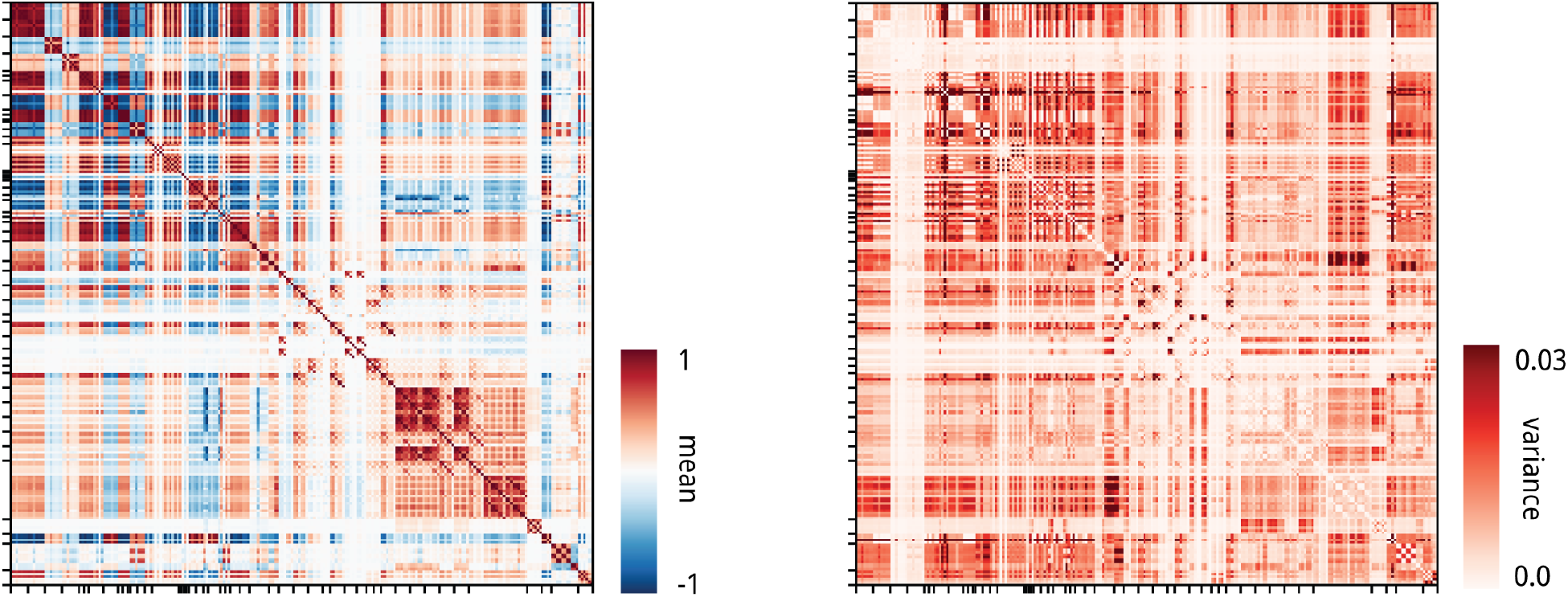
Similarity profile (Left) Group mean (same as in Fig. 1), and **(Right)** group variance matrices calculated from the individual similarity profiles across subjects and runs.

**Figure S2.**
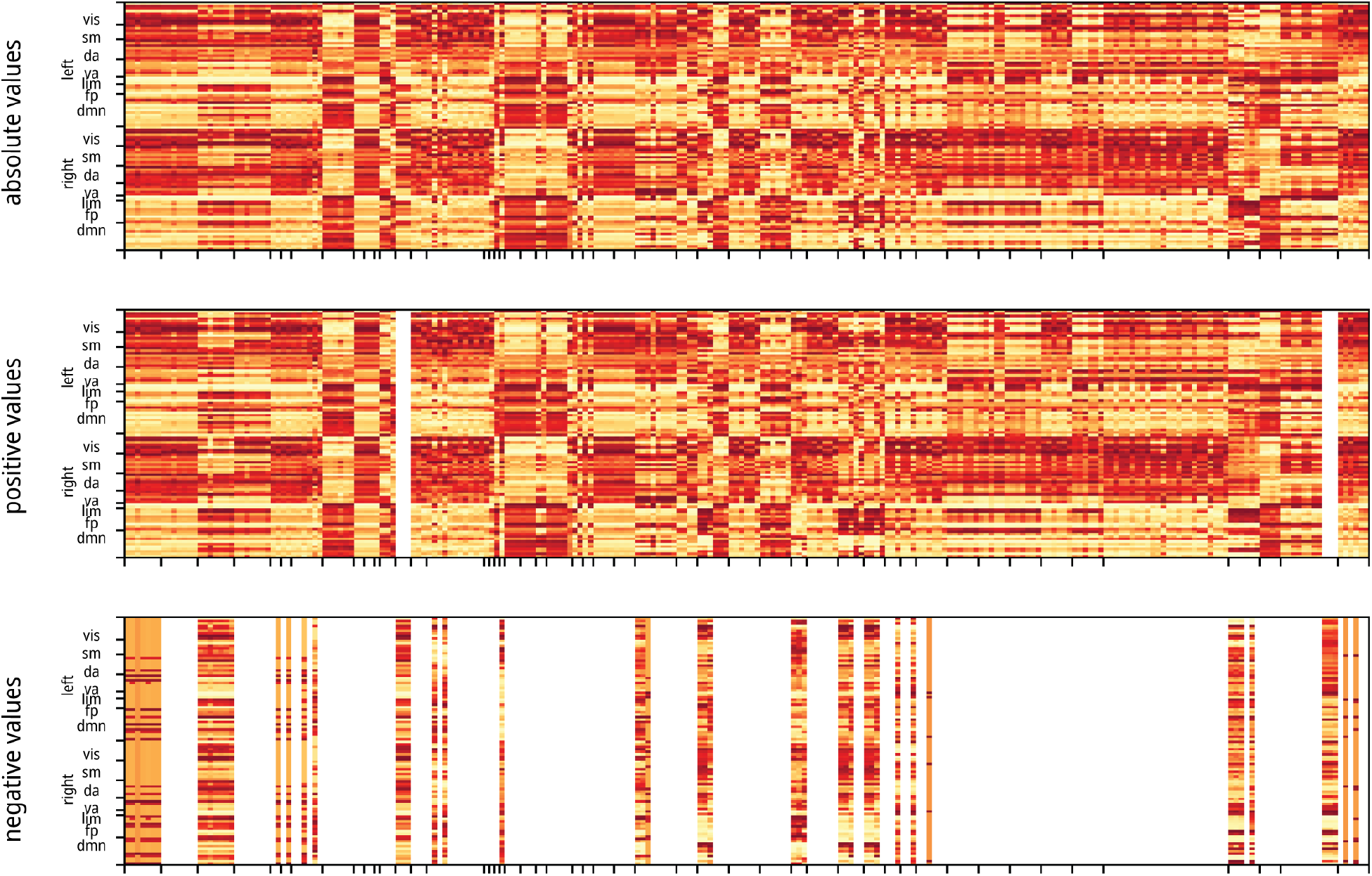
Hubness. Ranking of hubs quantified by weighted degree of the pairwise statistic matrices, shown for (from top to bottom) absolute values, positive values only, and negative values only. Rankings have not been not flipped like in Fig. 2b. Regions are ordered by intrinsic functional networks from [210] for left and right hemispheres. Darker red means more hubness. VIS: visual, SM: somatomotor, DA: dorsal attention, VA: ventral attention, LIM: limbic, FP: fronto-parietal, DMN: default mode network.

**Figure S3.**
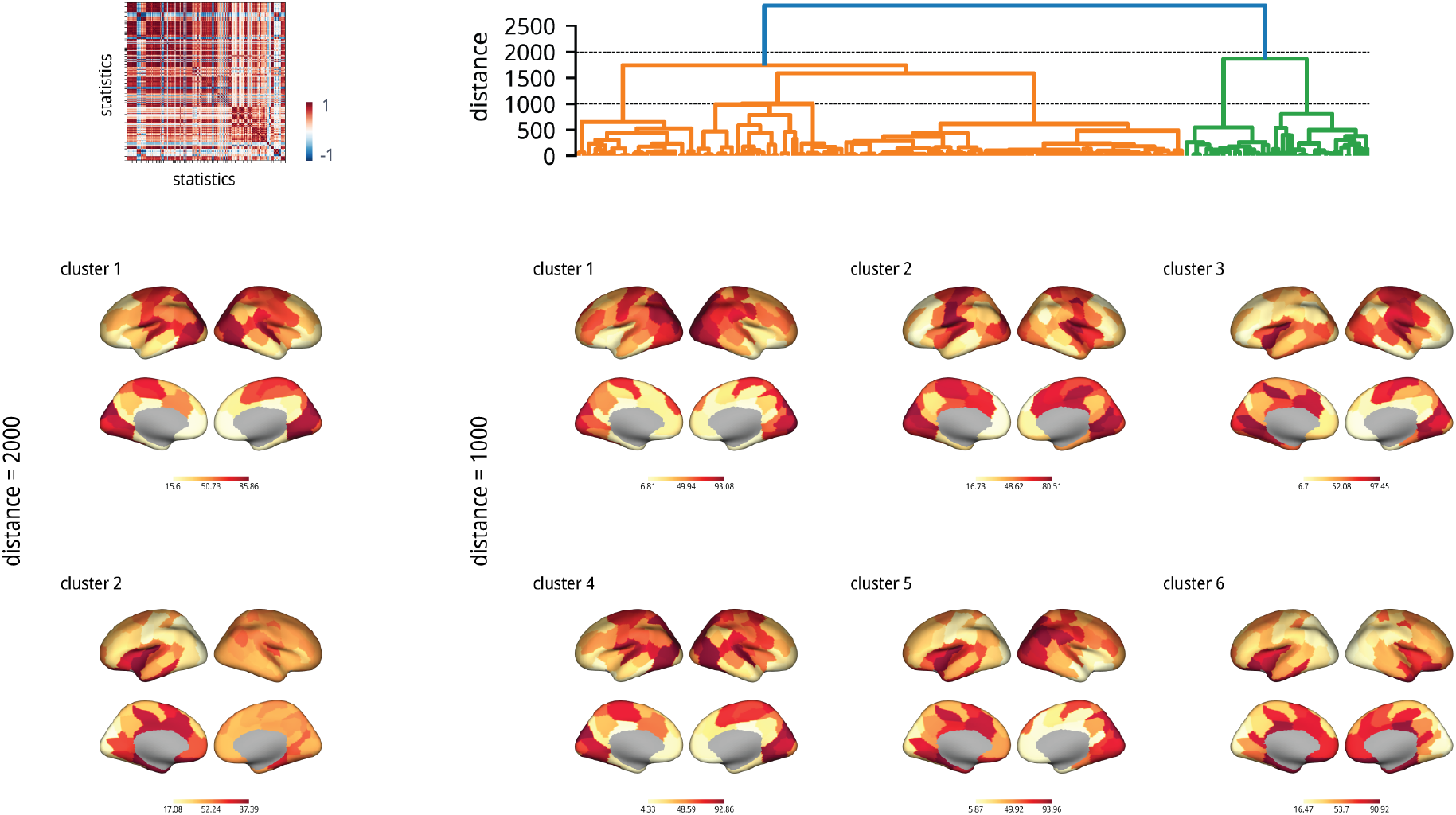
Hubness clusters. Hierarchical clustering of the hubness results in Fig. 2b. **(Upper left)** Similarity between the hubness results between the statistics in Fig. 2b. **(Upper right)** Dendrogram showing hierarchical clustering of the hubness results. **(Lower left)** The two clusters at *d* = 2000 averaged across statistics and shown on the cortical surface. **(Lower right)** The six clusters at *d* = 1000 averaged across statistics and shown on the cortical surface. Hierarchical clustering is implemented using *scipy* with *ward* linkage method.

**Figure S4.**
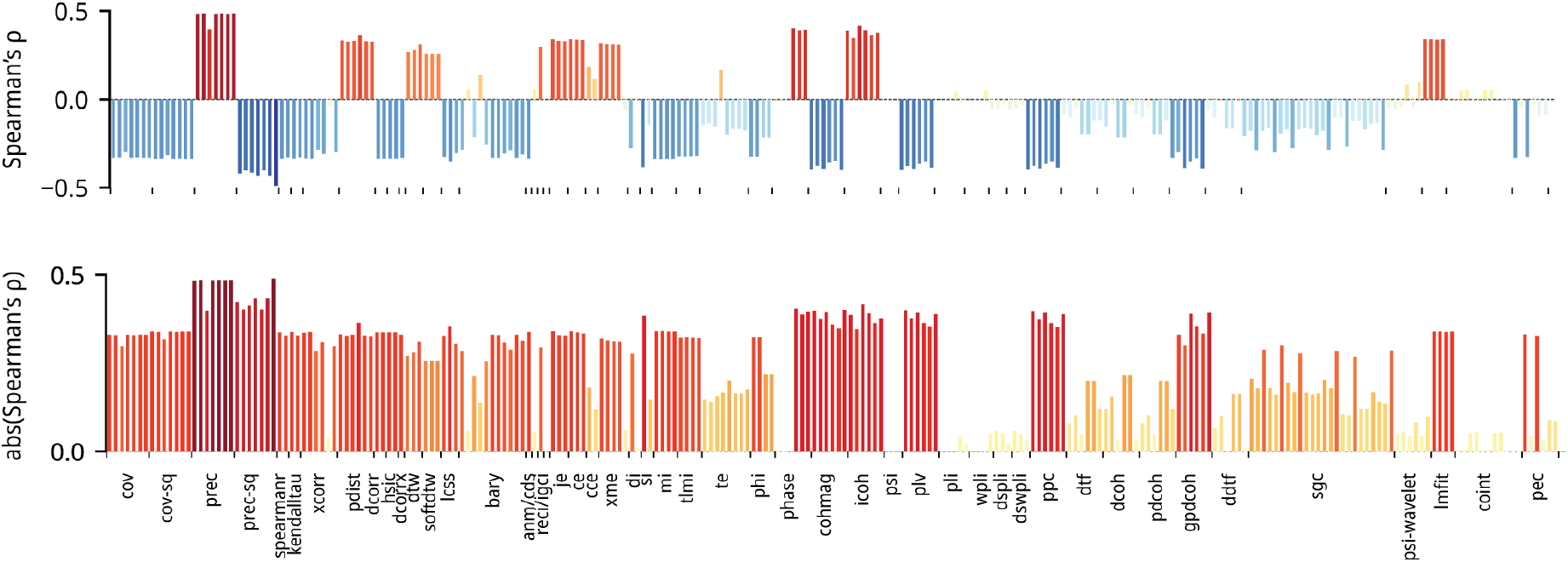
Structure-function coupling with Spearman’s rank correlation. To validate Fig. 2d, we calculated structure-function coupling as the Spearman’s rank correlation coefficient following Baum et al. [14]. The 2272 non-zero elements of the structural connectivity and corresponding values in the FC matrices were used to calculate the correlation.

**Figure S5.**
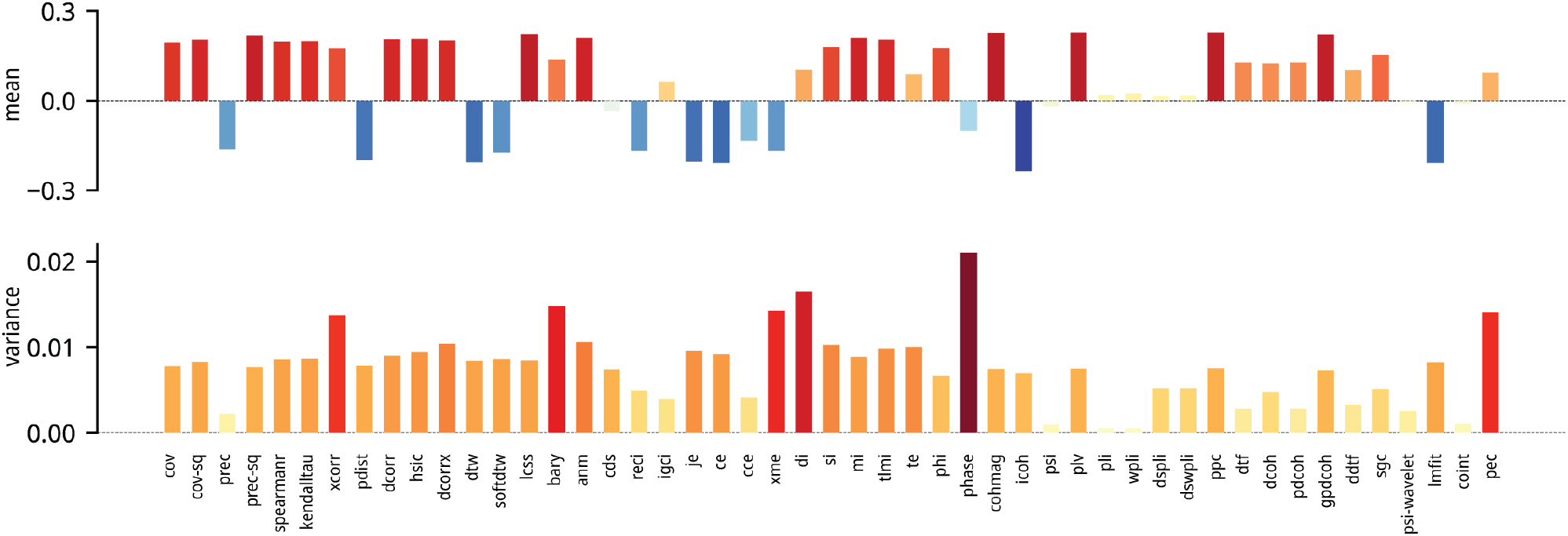
Multimodal neurophysiological networks. Correlations between statistics and the neurophysiological networks summarized across the five networks (correlated gene expression, laminar similarity, neurotransmitter receptor similarity, electrophysiological connectivity, and metabolic connectivity), showing **(Upper)** mean and **(Lower)** variance across the 49 measures.

**Figure S6.**
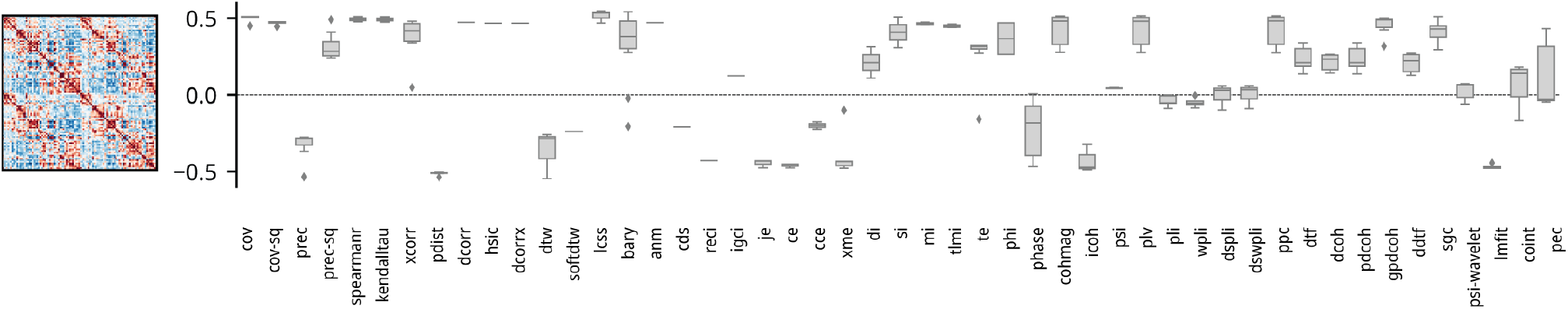
Correlation with cognitive similarity network. Similar to Fig. 3, pairwise interaction statistics matrices were correlated with inter-regional cognitive similarity network. Cognitive similarity network was estimated from regional functional associations of Neurosynth terms. Neurosynth is a meta-analytical database containing the voxel coordinates and related high-frequency keywords for >15,000 fMRI studies [209]. Briefly, we selected 123 neurocognitive terms from the Cognitive Atlas, a public ontology for cognitive science [137], and estimated a probabilistic measure of association for each term. The probabilistic measure can be interpreted as a quantitative account of regional activation relating to the neuropsychological process. A detailed methodological description can be found in Hansen et al. [65].

**Figure S7.**
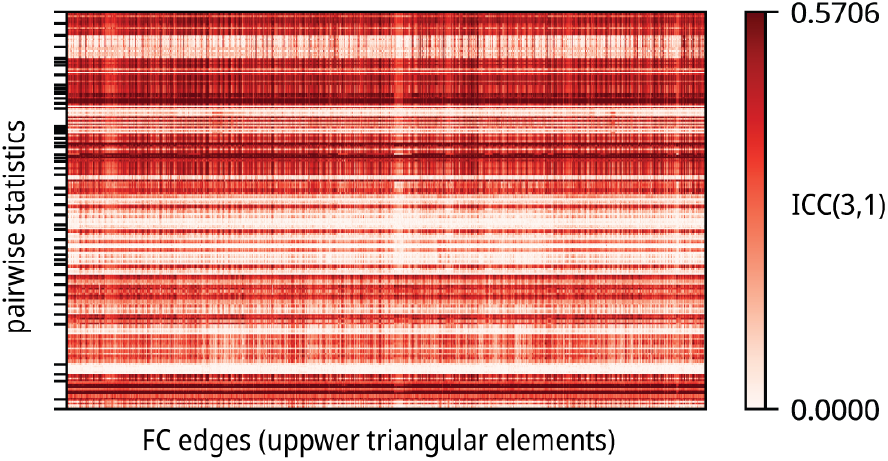
Intraclass correlation (ICC) for pairwise statistics. ICC was calculated for the pairwise statistics using the 4 restingstate runs for each participant. We used ICC(3, 1) implemented following *PyReliMRI* [40] and was defined in detail by Liljequist et al. [91].

**Figure S8.**
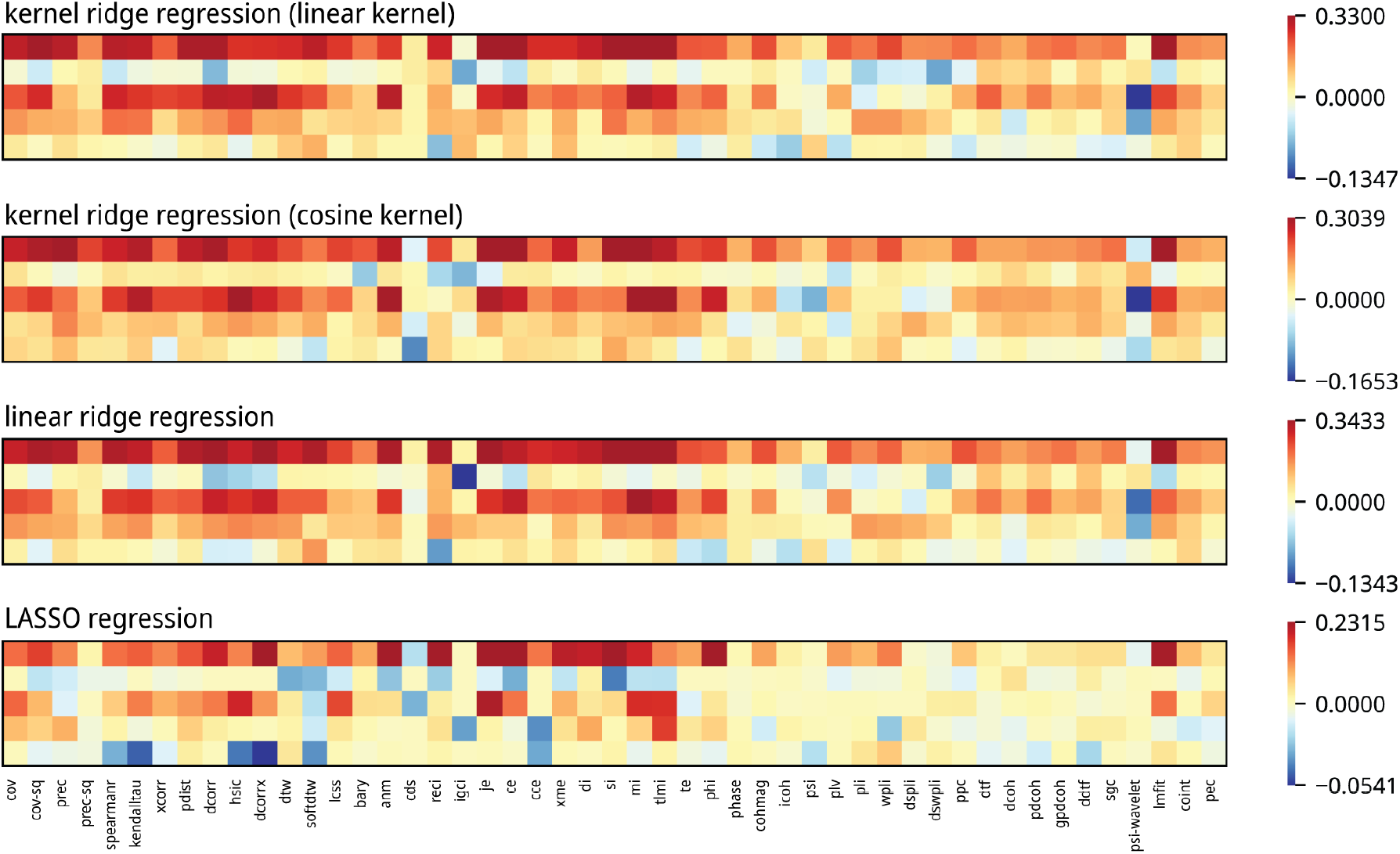
Behavior prediction. Mean prediction results from Fig. 4b (kernel ridge regression using linear kernel) are shown with alternative statistical learning algorithms, including kernel ridge regression (cosine kernel), linear ridge regression, and LASSO regression. The colorbars cover both negative values (0th percentile to 0; in blue) and positive values (0 to 97.5th percentile; in red).

**Figure S9.**
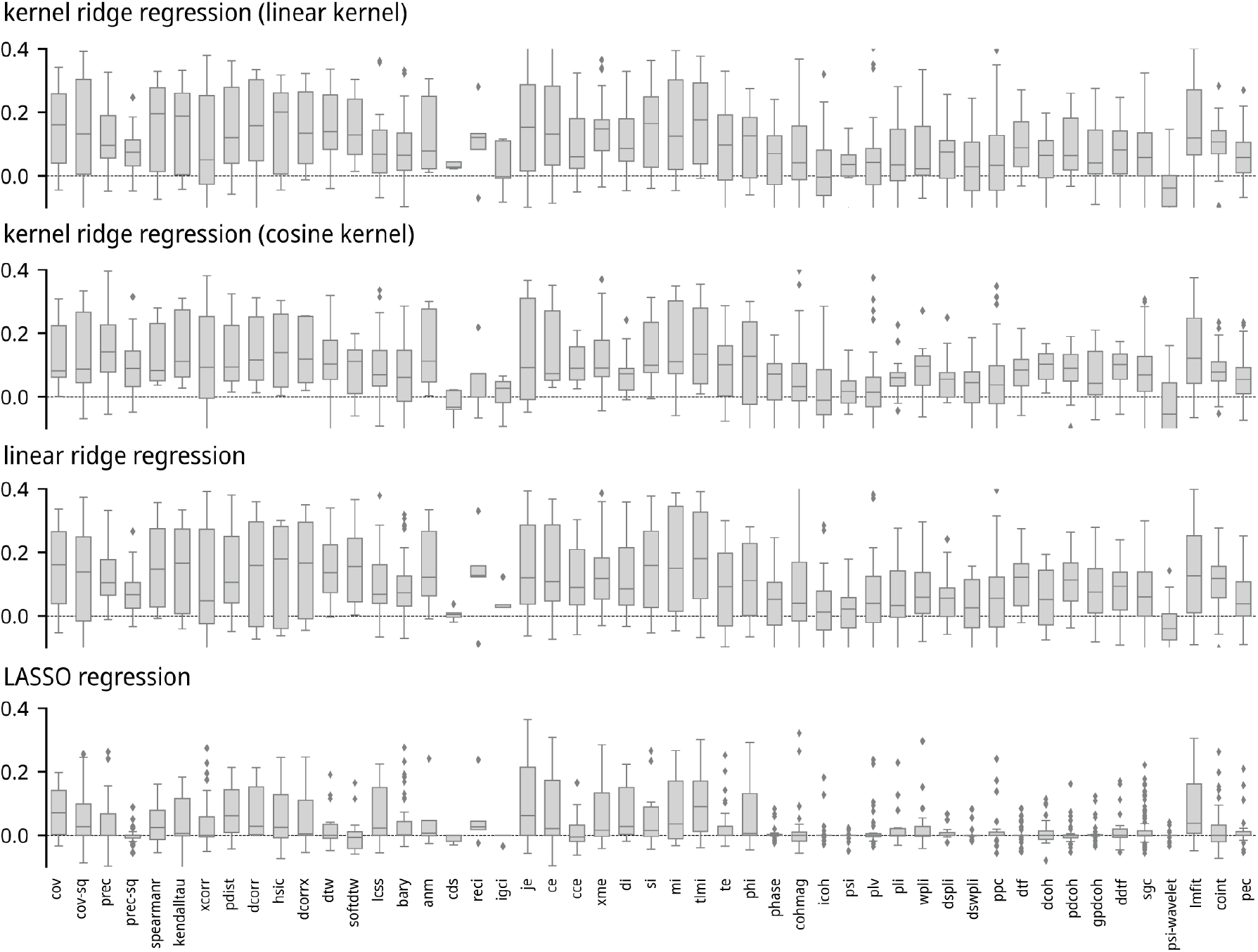
Behavior prediction. Prediction results from Fig. S8 are displayed in boxplots, showing individual statistic variations within each of the 49 measures.

**Figure S10.**
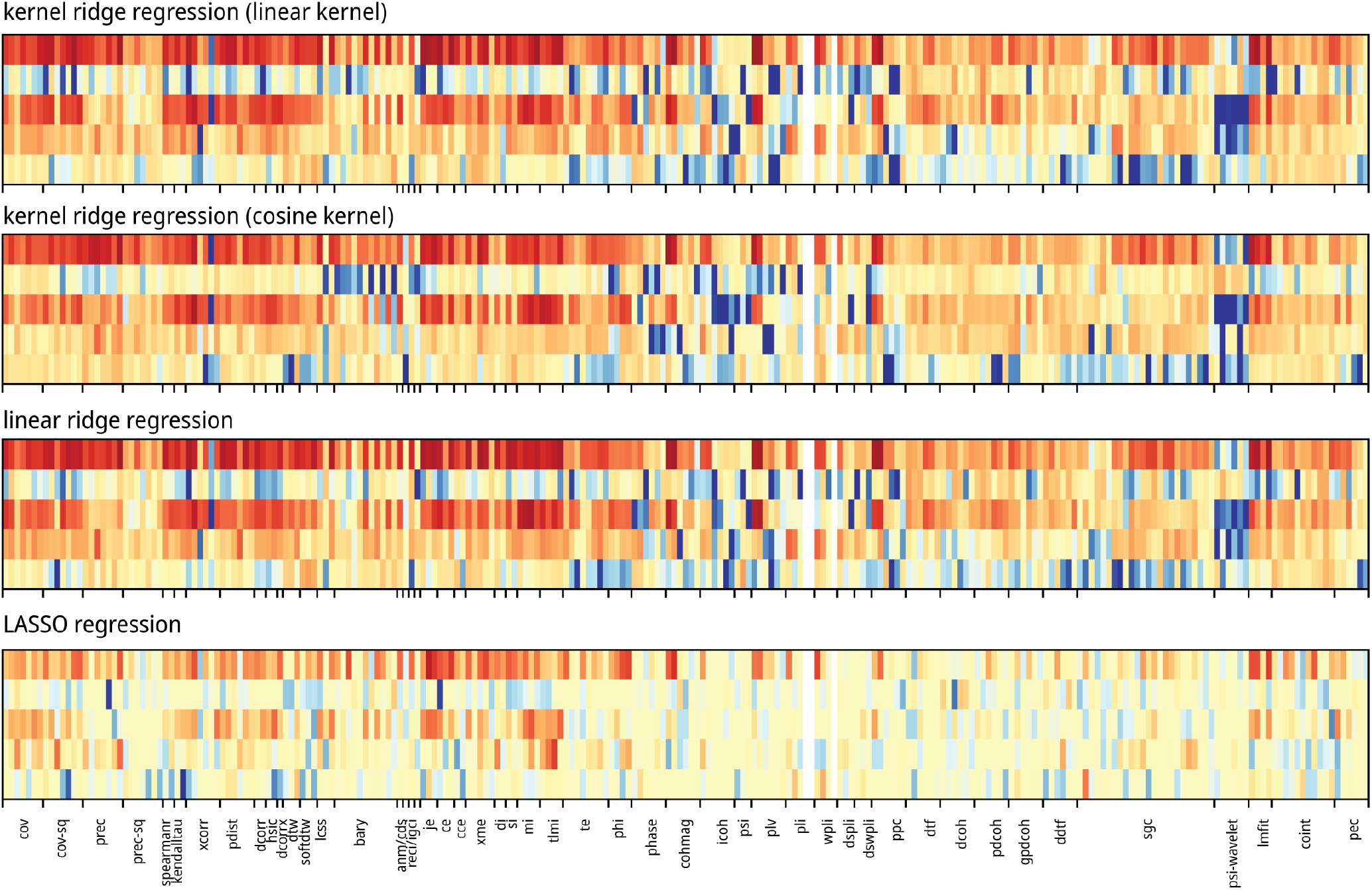
Behavior prediction. Mean prediction results from Fig. S8 are displayed for each of the 239 statistics.

**Figure S11.**
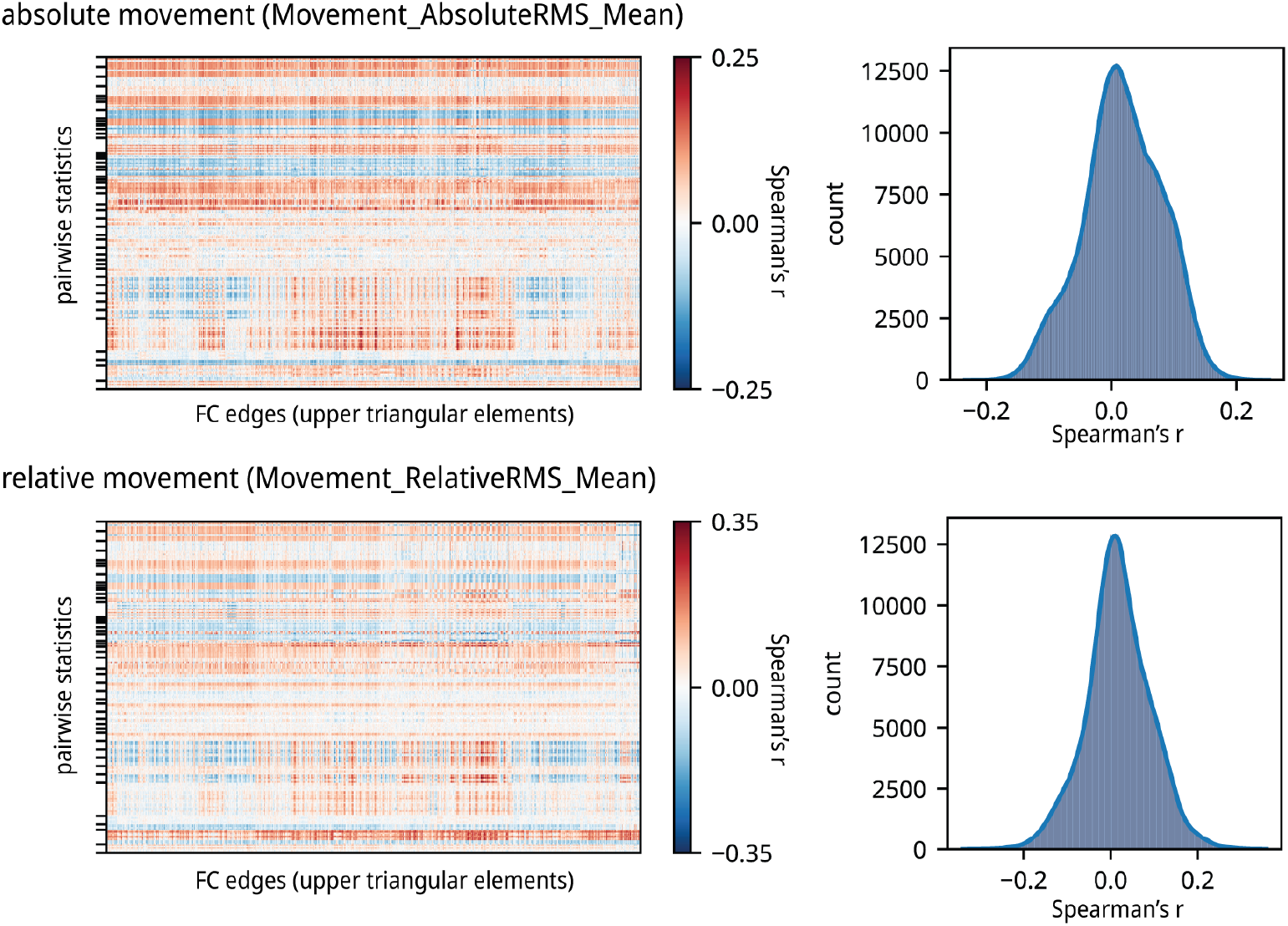
Sensitivity of pairwise statistics to participant movement. Root mean square of mean absolute (Upper) and relative (Lower) movement data during resting-state fMRI acquisition were correlated with each pairwise interaction statistic.

**Figure S12.**
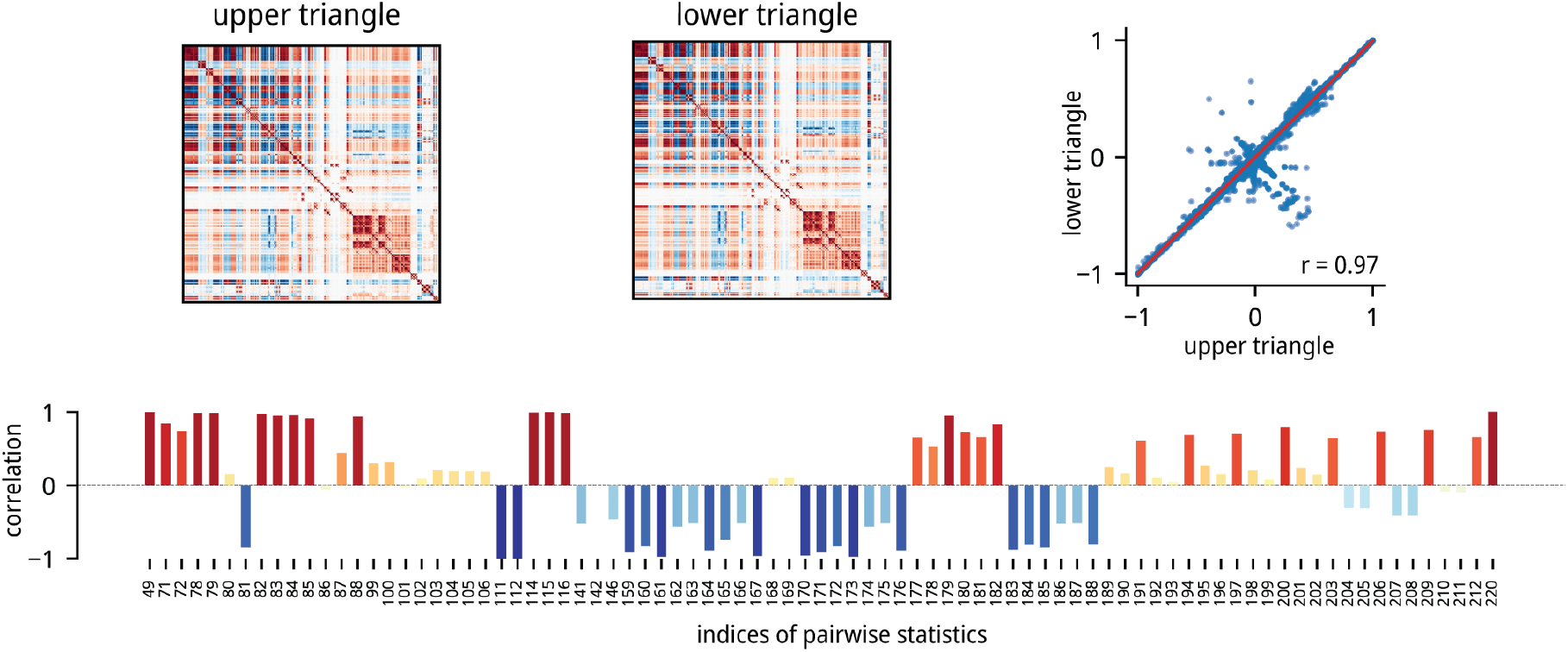
Directed measures in the pairwise statistics. We extracted the statistics with different upper and lower triangular values (excluding those which only differ by a sign) and study the implication of their directness. On the upper row, the group mean similarity profile (as in Fig. 1) was calculated using upper triangular values (used in the main results) and lower triangular values. Their correlation was calculated similar to Fig. 6. On the bottom row, Spearman’s rank correlations between the upper and lower triangular values were calculated for each directed pairwise statistics matrix. The indices on the x-axis correspond to the indices in Supplementary Table S1. A full list of directed pairwise statistics is shown in Supplementary Table S3. The list of pairwise statistics with only sign changes is shown in Supplementary Table S4

**TABLE S1:**
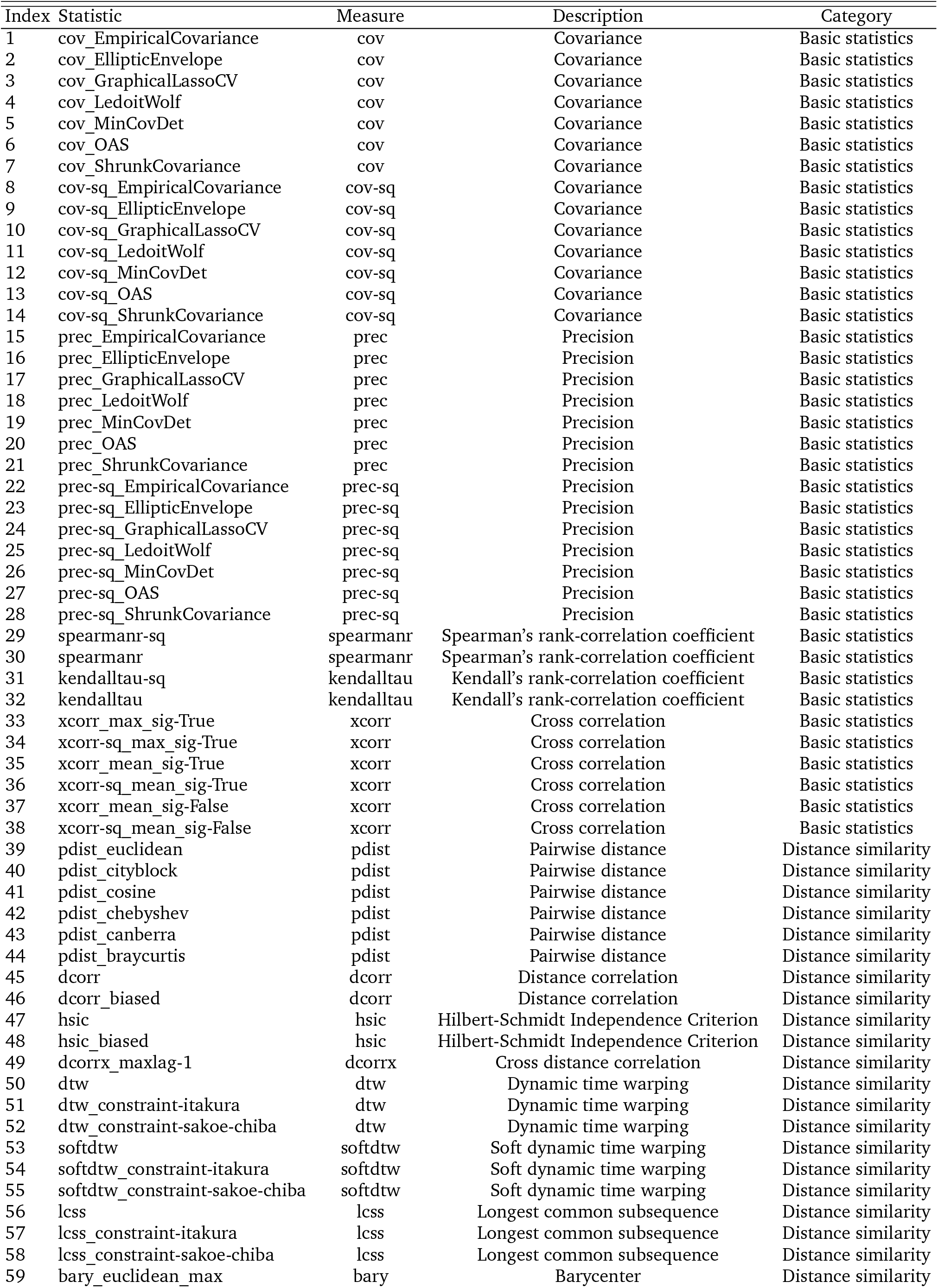
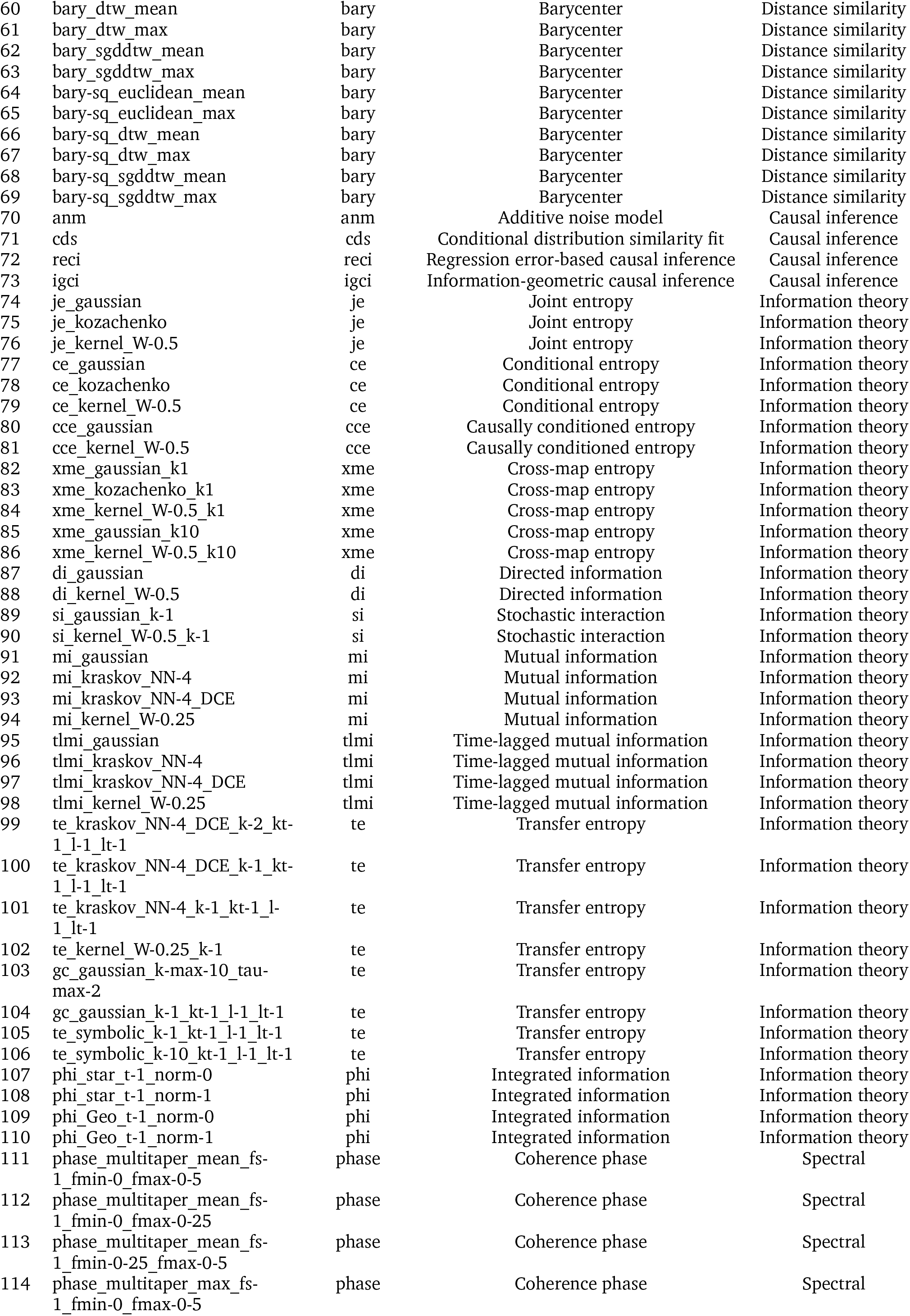
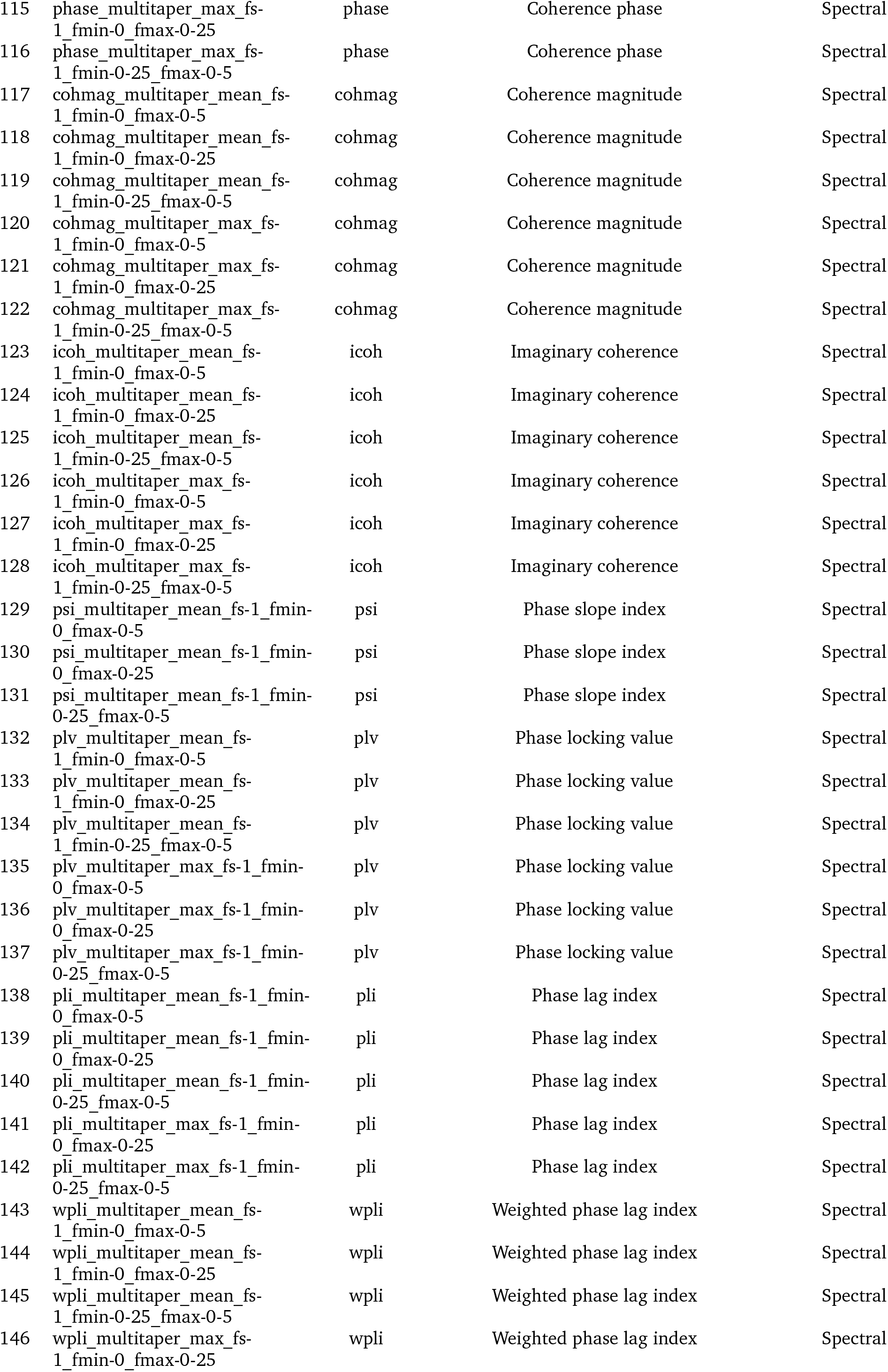
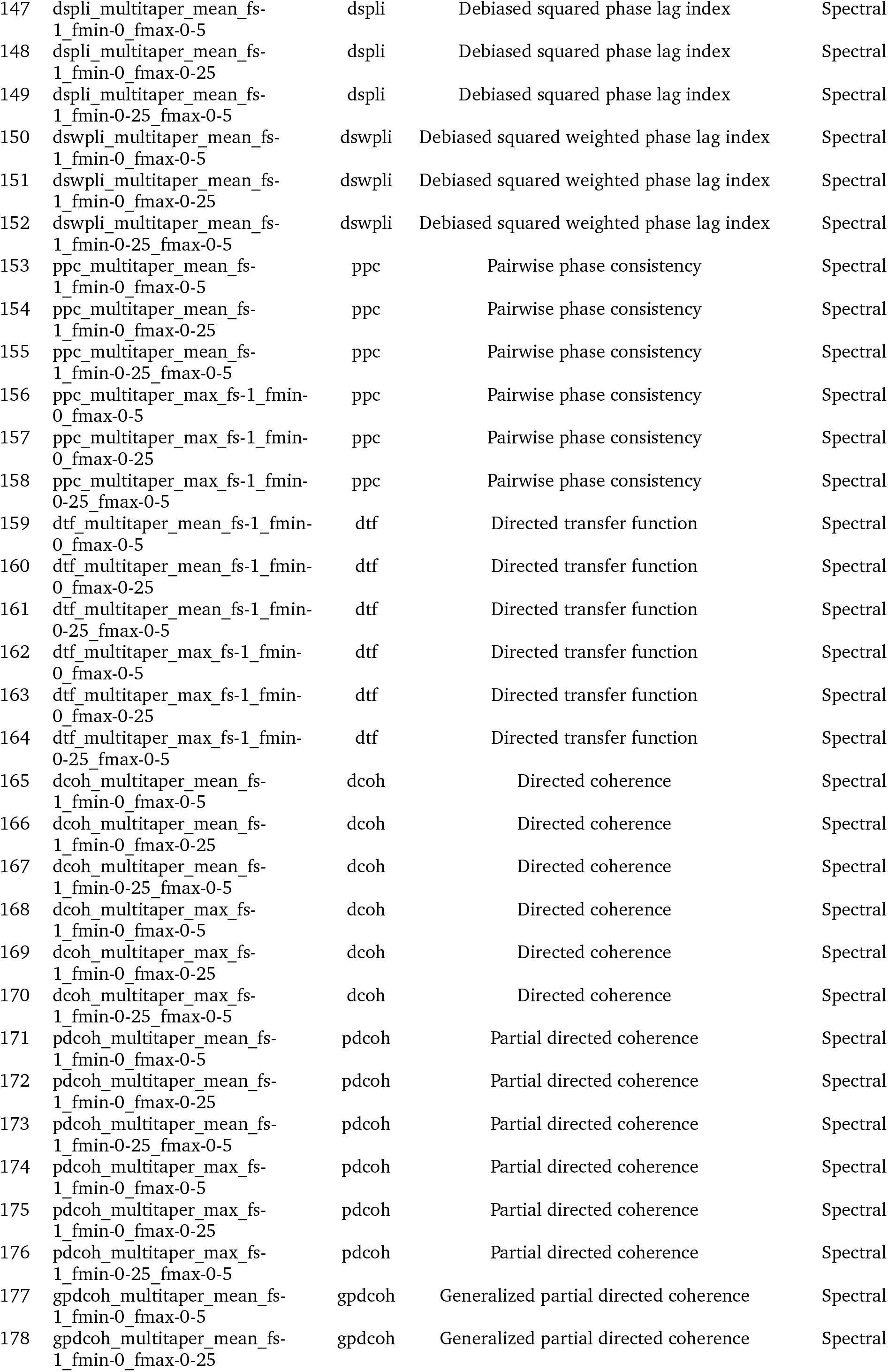
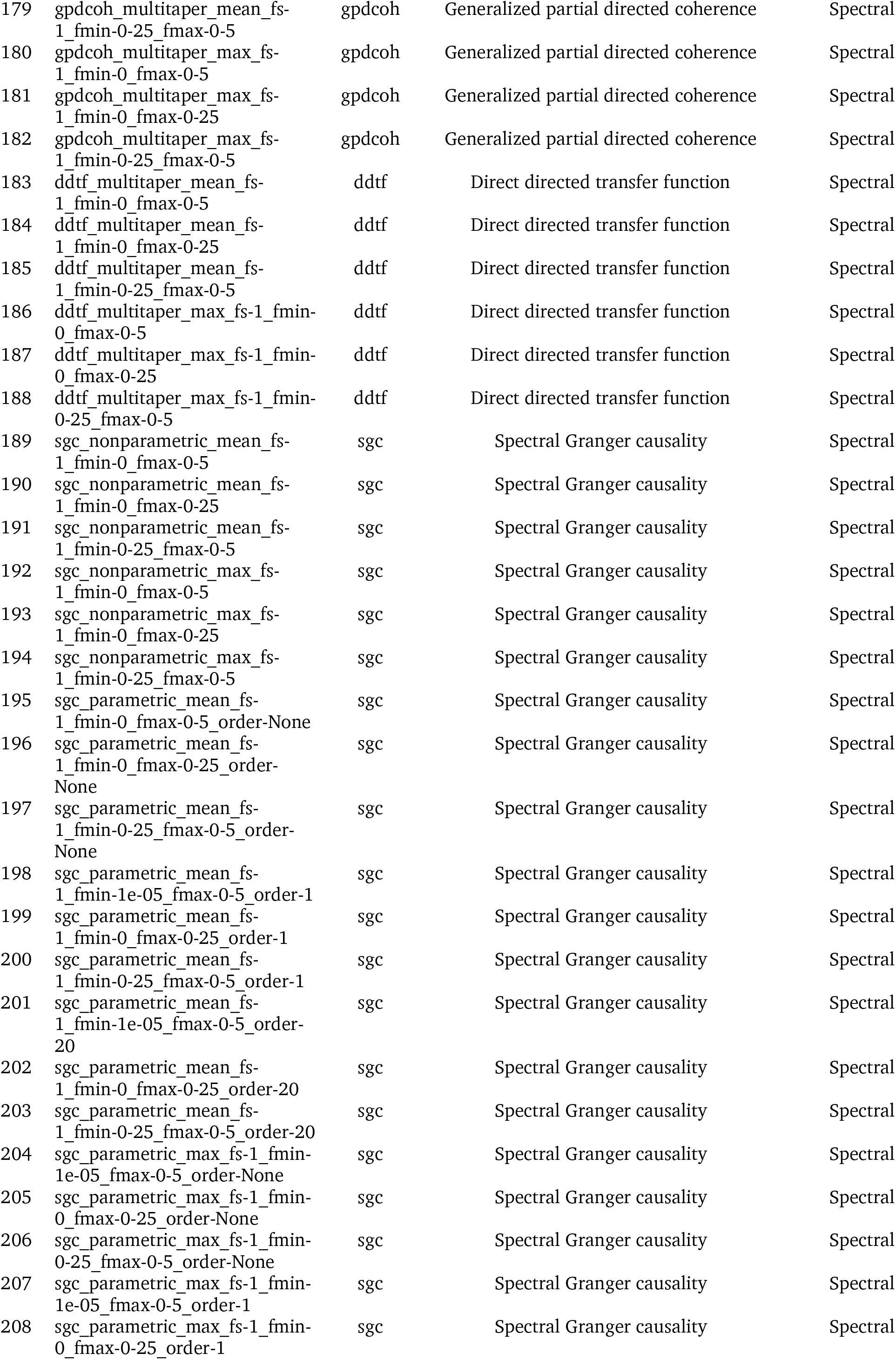
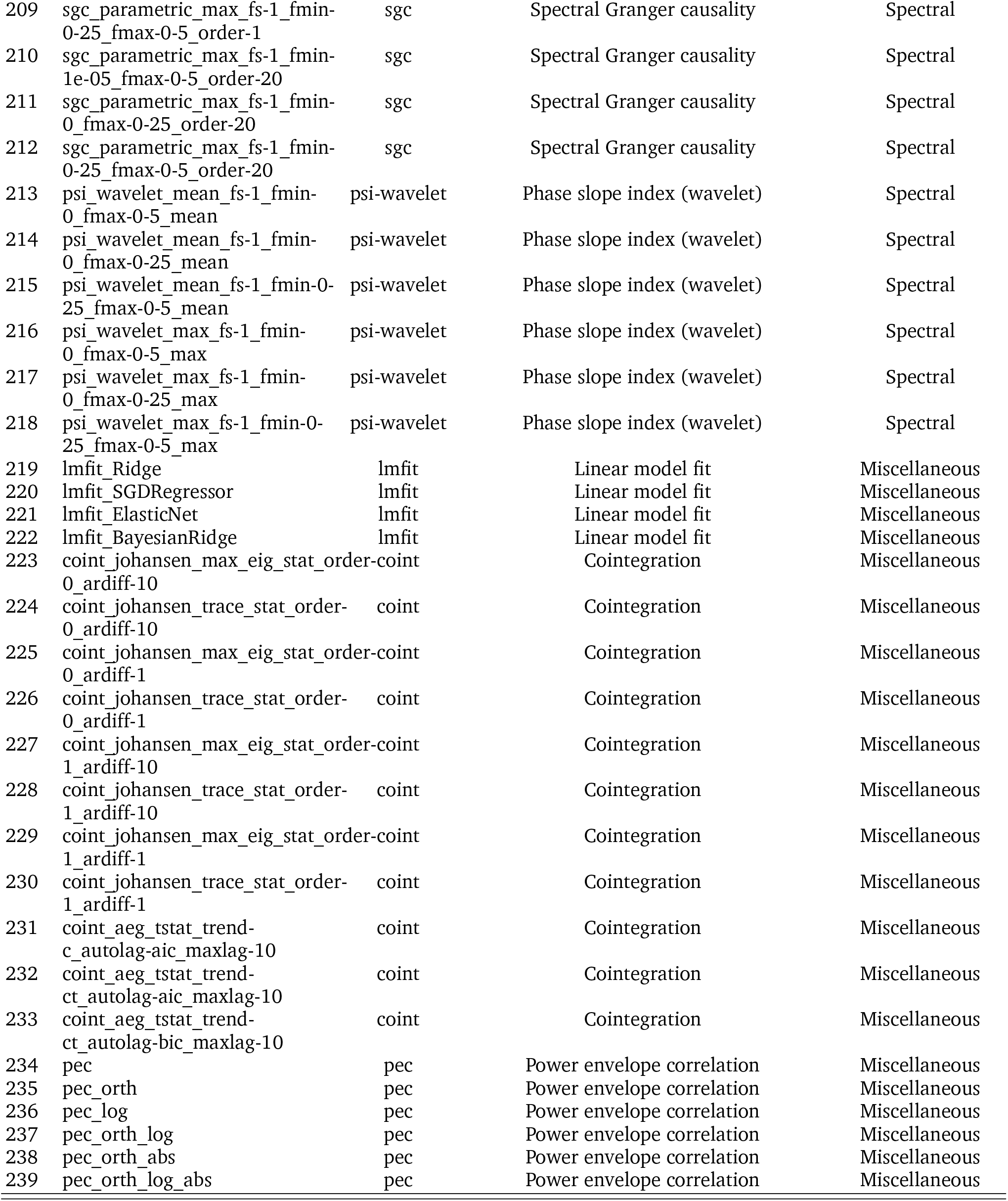
Full list of 239 *pyspi* statistics. The 239 pairwise statistics grouped into 49 measures across 6 major model families used in the main results.

**TABLE S2:** Reduced list of 179 *pyspi* statistics. The subset of 179 pairwise statistics used in the calculation of Fig. 6c.

| Index | Statistic | Measure | Description | Category |
| --- | --- | --- | --- | --- |
| 1 | cov_EmpiricalCovariance | cov | Covariance | Basic statistics |
| 2 | cov_EllipticEnvelope | cov | Covariance | Basic statistics |
| 3 | cov_GraphicalLassoCV | cov | Covariance | Basic statistics |
| 4 | cov_LedoitWolf | cov | Covariance | Basic statistics |
| 5 | cov_MinCovDet | cov | Covariance | Basic statistics |
| 6 | cov_OAS | cov | Covariance | Basic statistics |
| 7 | cov_ShrunkCovariance | cov | Covariance | Basic statistics |
| 8 | cov-sq_EmpiricalCovariance | cov-sq | Covariance | Basic statistics |
| 9 | cov-sq_EllipticEnvelope | cov-sq | Covariance | Basic statistics |
| 10 | cov-sq_GraphicalLassoCV | cov-sq | Covariance | Basic statistics |
| 11 | cov-sq_LedoitWolf | cov-sq | Covariance | Basic statistics |
| 12 | cov-sq_MinCovDet | cov-sq | Covariance | Basic statistics |
| 13 | cov-sq_OAS | cov-sq | Covariance | Basic statistics |
| 14 | cov-sq_ShrunkCovariance | cov-sq | Covariance | Basic statistics |
| 15 | prec_EmpiricalCovariance | prec | Precision | Basic statistics |
| 16 | prec_EllipticEnvelope | prec | Precision | Basic statistics |
| 17 | prec_GraphicalLassoCV | prec | Precision | Basic statistics |
| 18 | prec_LedoitWolf | prec | Precision | Basic statistics |
| 19 | prec_MinCovDet | prec | Precision | Basic statistics |
| 20 | prec_OAS | prec | Precision | Basic statistics |
| 21 | prec_ShrunkCovariance | prec | Precision | Basic statistics |
| 22 | prec-sq_EmpiricalCovariance | prec-sq | Precision | Basic statistics |
| 23 | prec-sq_EllipticEnvelope | prec-sq | Precision | Basic statistics |
| 24 | prec-sq_GraphicalLassoCV | prec-sq | Precision | Basic statistics |
| 25 | prec-sq_LedoitWolf | prec-sq | Precision | Basic statistics |
| 26 | prec-sq_MinCovDet | prec-sq | Precision | Basic statistics |
| 27 | prec-sq_OAS | prec-sq | Precision | Basic statistics |
| 28 | prec-sq_ShrunkCovariance | prec-sq | Precision | Basic statistics |
| 29 | spearmanr-sq | spearmanr | Spearman's rank-correlation coefficient | Basic statistics |
| 30 | spearmanr | spearmanr | Spearman's rank-correlation coefficient | Basic statistics |
| 31 | kendalltau-sq | kendalltau | Kendall's rank-correlation coefficient | Basic statistics |
| 32 | kendalltau | kendalltau | Kendall's rank-correlation coefficient | Basic statistics |
| 33 | xcorr_max_sig-True | xcorr | Cross correlation | Basic statistics |
| 34 | xcorr-sq_max_sig-True | xcorr | Cross correlation | Basic statistics |
| 35 | xcorr_mean_sig-True | xcorr | Cross correlation | Basic statistics |
| 36 | xcorr-sq_mean_sig-True | xcorr | Cross correlation | Basic statistics |
| 37 | xcorr_mean_sig-False | xcorr | Cross correlation | Basic statistics |
| 38 | xcorr-sq_mean_sig-False | xcorr | Cross correlation | Basic statistics |
| 39 | pdist_euclidean | pdist | Pairwise distance | Distance similarity |
| 40 | pdist_cityblock | pdist | Pairwise distance | Distance similarity |
| 41 | pdist_cosine | pdist | Pairwise distance | Distance similarity |
| 42 | pdist_chebyshev | pdist | Pairwise distance | Distance similarity |
| 43 | pdist_canberra | pdist | Pairwise distance | Distance similarity |
| 44 | pdist_braycurtis | pdist | Pairwise distance | Distance similarity |
| 45 | bary_euclidean_max | bary | Barycenter | Distance similarity |
| 46 | bary-sq_euclidean_mean | bary | Barycenter | Distance similarity |
| 47 | bary-sq_euclidean_max | bary | Barycenter | Distance similarity |
| 48 | je_gaussian | je | Joint entropy | Information theory |
| 49 | je_kernel_W-0.5 | je | Joint entropy | Information theory |
| 50 | ce_gaussian | ce | Conditional entropy | Information theory |
| 51 | ce_kozachenko | ce | Conditional entropy | Information theory |
| 52 | ce_kernel_W-0.5 | ce | Conditional entropy | Information theory |
| 53 | xme_gaussian_k1 | xme | Cross-map entropy | Information theory |
| 54 | xme_gaussian_k10 | xme | Cross-map entropy | Information theory |
| 55 | xme_kernel_W-0.5_k10 | xme | Cross-map entropy | Information theory |
| 56 | mi_gaussian | mi | Mutual information | Information theory |
| 57 | tlmi_gaussian | tlmi | Time-lagged mutual information | Information theory |
| 58 | tlmi_kraskov_NN-4 | tlmi | Time-lagged mutual information | Information theory |
| 59 | tlmi_kraskov_NN-4_DCE | tlmi | Time-lagged mutual information | Information theory |
| 60 | gc_gaussian_k-1_kt-1_l-1_lt-1 | te | Transfer entropy | Information theory |
| 61 | te_symbolic_k-1_kt-1_l-1_lt-1 | te | Transfer entropy | Information theory |
| 62 | te_symbolic_k-10_kt-1_l-1_lt-1 | te | Transfer entropy | Information theory |
| 63 | phase_multitaper_mean_fs-1_fmin-0_fmax-0-5 | phase | Coherence phase | Spectral |
| 64 | phase_multitaper_mean_fs-1_fmin-0_fmax-0-25 | phase | Coherence phase | Spectral |
| 65 | phase_multitaper_mean_fs-1_fmin-0-25_fmax-0-5 | phase | Coherence phase | Spectral |
| 66 | phase_multitaper_max_fs-1_fmin-0_fmax-0-5 | phase | Coherence phase | Spectral |
| 67 | phase_multitaper_max_fs-1_fmin-0_fmax-0-25 | phase | Coherence phase | Spectral |
| 68 | phase_multitaper_max_fs-1_fmin-0-25_fmax-0-5 | phase | Coherence phase | Spectral |
| 69 | cohmag_multitaper_mean_fs-1_fmin-0_fmax-0-5 | cohmag | Coherence magnitude | Spectral |
| 70 | cohmag_multitaper_mean_fs-1_fmin-0_fmax-0-25 | cohmag | Coherence magnitude | Spectral |
| 71 | cohmag_multitaper_mean_fs-1_fmin-0-25_fmax-0-5 | cohmag | Coherence magnitude | Spectral |
| 72 | cohmag_multitaper_max_fs-1_fmin-0_fmax-0-5 | cohmag | Coherence magnitude | Spectral |
| 73 | cohmag_multitaper_max_fs-1_fmin-0_fmax-0-25 | cohmag | Coherence magnitude | Spectral |
| 74 | cohmag_multitaper_max_fs-1_fmin-0-25_fmax-0-5 | cohmag | Coherence magnitude | Spectral |
| 75 | icoh_multitaper_mean_fs-1_fmin-0_fmax-0-5 | icoh | Imaginary coherence | Spectral |
| 76 | icoh_multitaper_mean_fs-1_fmin-0_fmax-0-25 | icoh | Imaginary coherence | Spectral |
| 77 | icoh_multitaper_mean_fs-1_fmin-0-25_fmax-0-5 | icoh | Imaginary coherence | Spectral |
| 78 | icoh_multitaper_max_fs-1_fmin-0_fmax-0-5 | icoh | Imaginary coherence | Spectral |
| 79 | icoh_multitaper_max_fs-1_fmin-0_fmax-0-25 | icoh | Imaginary coherence | Spectral |
| 80 | icoh_multitaper_max_fs-1_fmin-0-25_fmax-0-5 | icoh | Imaginary coherence | Spectral |
| 81 | psi_multitaper_mean_fs-1_fmin-0_fmax-0-5 | psi | Phase slope index | Spectral |
| 82 | psi_multitaper_mean_fs-1_fmin-0_fmax-0-25 | psi | Phase slope index | Spectral |
| 83 | psi_multitaper_mean_fs-1_fmin-0-25_fmax-0-5 | psi | Phase slope index | Spectral |
| 84 | pli_multitaper_mean_fs-1_fmin-0_fmax-0-5 | pli | Phase lag index | Spectral |
| 85 | pli_multitaper_mean_fs-1_fmin-0_fmax-0-25 | pli | Phase lag index | Spectral |
| 86 | pli_multitaper_mean_fs-1_fmin-0-25_fmax-0-5 | pli | Phase lag index | Spectral |
| 87 | pli_multitaper_max_fs-1_fmin-0_fmax-0-25 | pli | Phase lag index | Spectral |
| 88 | pli_multitaper_max_fs-1_fmin-0-25_fmax-0-5 | pli | Phase lag index | Spectral |
| 89 | wpli_multitaper_mean_fs-1_fmin-0_fmax-0-5 | wpli | Weighted phase lag index | Spectral |
| 90 | wpli_multitaper_mean_fs-1_fmin-0_fmax-0-25 | wpli | Weighted phase lag index | Spectral |
| 91 | wpli_multitaper_mean_fs-1_fmin-0-25_fmax-0-5 | wpli | Weighted phase lag index | Spectral |
| 92 | wpli_multitaper_max_fs-1_fmin-0-25_fmax-0-5 | wpli | Weighted phase lag index | Spectral |
| 93 | dspli_multitaper_mean_fs-1_fmin-0_fmax-0-5 | dspli | Debiased squared phase lag index | Spectral |
| 94 | dspli_multitaper_mean_fs-1_fmin-0_fmax-0-25 | dspli | Debiased squared phase lag index | Spectral |
| 95 | dspli_multitaper_mean_fs-1_fmin-0-25_fmax-0-5 | dspli | Debiased squared phase lag index | Spectral |
| 96 | dswpli_multitaper_mean_fs-1_fmin-0_fmax-0-5 | dswpli | Debiased squared weighted phase lag index | Spectral |
| 97 | dswpli_multitaper_mean_fs-1_fmin-0_fmax-0-25 | dswpli | Debiased squared weighted phase lag index | Spectral |
| 98 | dswpli_multitaper_mean_fs-1_fmin-0-25_fmax-0-5 | dswpli | Debiased squared weighted phase lag index | Spectral |
| 99 | ppc_multitaper_mean_fs-1_fmin-0_fmax-0-5 | ppc | Pairwise phase consistency | Spectral |
| 100 | ppc_multitaper_mean_fs-1_fmin-0_fmax-0-25 | ppc | Pairwise phase consistency | Spectral |
| 101 | ppc_multitaper_mean_fs-1_fmin-0-25_fmax-0-5 | ppc | Pairwise phase consistency | Spectral |
| 102 | ppc_multitaper_max_fs-1_fmin-0_fmax-0-5 | ppc | Pairwise phase consistency | Spectral |
| 103 | ppc_multitaper_max_fs-1_fmin-0_fmax-0-25 | ppc | Pairwise phase consistency | Spectral |
| 104 | ppc_multitaper_max_fs-1_fmin-0-25_fmax-0-5 | ppc | Pairwise phase consistency | Spectral |
| 105 | dtf_multitaper_mean_fs-1_fmin-0_fmax-0-5 | dtf | Directed transfer function | Spectral |
| 106 | dtf_multitaper_mean_fs-1_fmin-0_fmax-0-25 | dtf | Directed transfer function | Spectral |
| 107 | dtf_multitaper_mean_fs-1_fmin-0-25_fmax-0-5 | dtf | Directed transfer function | Spectral |
| 108 | dtf_multitaper_max_fs-1_fmin-0_fmax-0-5 | dtf | Directed transfer function | Spectral |
| 109 | dtf_multitaper_max_fs-1_fmin-0_fmax-0-25 | dtf | Directed transfer function | Spectral |
| 110 | dtf_multitaper_max_fs-1_fmin-0-25_fmax-0-5 | dtf | Directed transfer function | Spectral |
| 111 | dcoh_multitaper_mean_fs-1_fmin-0_fmax-0-5 | dcoh | Directed coherence | Spectral |
| 112 | dcoh_multitaper_mean_fs-1_fmin-0_fmax-0-25 | dcoh | Directed coherence | Spectral |
| 113 | dcoh_multitaper_mean_fs-1_fmin-0-25_fmax-0-5 | dcoh | Directed coherence | Spectral |
| 114 | dcoh_multitaper_max_fs-1_fmin-0_fmax-0-5 | dcoh | Directed coherence | Spectral |
| 115 | dcoh_multitaper_max_fs-1_fmin-0_fmax-0-25 | dcoh | Directed coherence | Spectral |
| 116 | dcoh_multitaper_max_fs-1_fmin-0-25_fmax-0-5 | dcoh | Directed coherence | Spectral |
| 117 | pdcoh_multitaper_mean_fs-1_fmin-0_fmax-0-5 | pdcoh | Partial directed coherence | Spectral |
| 118 | pdcoh_multitaper_mean_fs-1_fmin-0_fmax-0-25 | pdcoh | Partial directed coherence | Spectral |
| 119 | pdcoh_multitaper_mean_fs-1_fmin-0-25_fmax-0-5 | pdcoh | Partial directed coherence | Spectral |
| 120 | pdcoh_multitaper_max_fs-1_fmin-0_fmax-0-5 | pdcoh | Partial directed coherence | Spectral |
| 121 | pdcoh_multitaper_max_fs-1_fmin-0_fmax-0-25 | pdcoh | Partial directed coherence | Spectral |
| 122 | pdcoh_multitaper_max_fs-1_fmin-0-25_fmax-0-5 | pdcoh | Partial directed coherence | Spectral |
| 123 | gpdcoh_multitaper_mean_fs-1_fmin-0_fmax-0-5 | gpdcoh | Generalized partial directed coherence | Spectral |
| 124 | gpdcoh_multitaper_mean_fs-1_fmin-0_fmax-0-25 | gpdcoh | Generalized partial directed coherence | Spectral |
| 125 | gpdcoh_multitaper_mean_fs-1_fmin-0-25_fmax-0-5 | gpdcoh | Generalized partial directed coherence | Spectral |
| 126 | gpdcoh_multitaper_max_fs-1_fmin-0_fmax-0-5 | gpdcoh | Generalized partial directed coherence | Spectral |
| 127 | gpdcoh_multitaper_max_fs-1_fmin-0_fmax-0-25 | gpdcoh | Generalized partial directed coherence | Spectral |
| 128 | gpdcoh_multitaper_max_fs-1_fmin-0-25_fmax-0-5 | gpdcoh | Generalized partial directed coherence | Spectral |
| 129 | ddtf_multitaper_mean_fs-1_fmin-0_fmax-0-5 | ddtf | Direct directed transfer function | Spectral |
| 130 | ddtf_multitaper_mean_fs-1_fmin-0_fmax-0-25 | ddtf | Direct directed transfer function | Spectral |
| 131 | ddtf_multitaper_mean_fs-1_fmin-0-25_fmax-0-5 | ddtf | Direct directed transfer function | Spectral |
| 132 | ddtf_multitaper_max_fs-1_fmin-0_fmax-0-5 | ddtf | Direct directed transfer function | Spectral |
| 133 | ddtf_multitaper_max_fs-1_fmin-0_fmax-0-25 | ddtf | Direct directed transfer function | Spectral |
| 134 | ddtf_multitaper_max_fs-1_fmin-0-25_fmax-0-5 | ddtf | Direct directed transfer function | Spectral |
| 135 | sgc_nonparametric_mean_fs-1_fmin-0_fmax-0-5 | sgc | Spectral Granger causality | Spectral |
| 136 | sgc_nonparametric_mean_fs-1_fmin-0_fmax-0-25 | sgc | Spectral Granger causality | Spectral |
| 137 | sgc_nonparametric_mean_fs-1_fmin-0-25_fmax-0-5 | sgc | Spectral Granger causality | Spectral |
| 138 | sgc_nonparametric_max_fs-1_fmin-0_fmax-0-5 | sgc | Spectral Granger causality | Spectral |
| 139 | sgc_nonparametric_max_fs-1_fmin-0_fmax-0-25 | sgc | Spectral Granger causality | Spectral |
| 140 | sgc_nonparametric_max_fs-1_fmin-0-25_fmax-0-5 | sgc | Spectral Granger causality | Spectral |
| 141 | sgc_parametric_mean_fs-1_fmin-0_fmax-0-5_order-None | sgc | Spectral Granger causality | Spectral |
| 142 | sgc_parametric_mean_fs-1_fmin-0_fmax-0-25_order-None | sgc | Spectral Granger causality | Spectral |
| 143 | sgc_parametric_mean_fs-1_fmin-0-25_fmax-0-5_order-None | sgc | Spectral Granger causality | Spectral |
| 144 | sgc_parametric_mean_fs-1_fmin-1e-05_fmax-0-5_order-1 | sgc | Spectral Granger causality | Spectral |
| 145 | sgc_parametric_mean_fs-1_fmin-0_fmax-0-25_order-1 | sgc | Spectral Granger causality | Spectral |
| 146 | sgc_parametric_mean_fs-1_fmin-0-25_fmax-0-5_order-1 | sgc | Spectral Granger causality | Spectral |
| 147 | sgc_parametric_mean_fs-1_fmin-1e-05_fmax-0-5_order-20 | sgc | Spectral Granger causality | Spectral |
| 148 | sgc_parametric_mean_fs-1_fmin-0_fmax-0-25_order-20 | sgc | Spectral Granger causality | Spectral |
| 149 | sgc_parametric_mean_fs-1_fmin-0-25_fmax-0-5_order-20 | sgc | Spectral Granger causality | Spectral |
| 150 | sgc_parametric_max_fs-1_fmin-1e-05_fmax-0-5_order-None | sgc | Spectral Granger causality | Spectral |
| 151 | sgc_parametric_max_fs-1_fmin-0_fmax-0-25_order-None | sgc | Spectral Granger causality | Spectral |
| 152 | sgc_parametric_max_fs-1_fmin-0-25_fmax-0-5_order-None | sgc | Spectral Granger causality | Spectral |
| 153 | sgc_parametric_max_fs-1_fmin-1e-05_fmax-0-5_order-1 | sgc | Spectral Granger causality | Spectral |
| 154 | sgc_parametric_max_fs-1_fmin-0_fmax-0-25_order-1 | sgc | Spectral Granger causality | Spectral |
| 155 | sgc_parametric_max_fs-1_fmin-0-25_fmax-0-5_order-1 | sgc | Spectral Granger causality | Spectral |
| 156 | sgc_parametric_max_fs-1_fmin-1e-05_fmax-0-5_order-20 | sgc | Spectral Granger causality | Spectral |
| 157 | sgc_parametric_max_fs-1_fmin-0_fmax-0-25_order-20 | sgc | Spectral Granger causality | Spectral |
| 158 | sgc_parametric_max_fs-1_fmin-0-25_fmax-0-5_order-20 | sgc | Spectral Granger causality | Spectral |
| 159 | lmfit_Ridge | lmfit | Linear model fit | Miscellaneous |
| 160 | lmfit_SGDRegressor | lmfit | Linear model fit | Miscellaneous |
| 161 | lmfit_ElasticNet | lmfit | Linear model fit | Miscellaneous |
| 162 | lmfit_BayesianRidge | lmfit | Linear model fit | Miscellaneous |
| 163 | coint_johansen_max_eig_stat_order-0_ardiff-10 | coint | Cointegration | Miscellaneous |
| 164 | coint_johansen_trace_stat_order-0_ardiff-10 | coint | Cointegration | Miscellaneous |
| 165 | coint_johansen_max_eig_stat_order-0_ardiff-1 | coint | Cointegration | Miscellaneous |
| 166 | coint_johansen_trace_stat_order-0_ardiff-1 | coint | Cointegration | Miscellaneous |
| 167 | coint_johansen_max_eig_stat_order-1_ardiff-10 | coint | Cointegration | Miscellaneous |
| 168 | coint_johansen_trace_stat_order-1_ardiff-10 | coint | Cointegration | Miscellaneous |
| 169 | coint_johansen_max_eig_stat_order-1_ardiff-1 | coint | Cointegration | Miscellaneous |
| 170 | coint_johansen_trace_stat_order-1_ardiff-1 | coint | Cointegration | Miscellaneous |
| 171 | coint_aeg_tstat_trend-c_autolag-aic_maxlag-10 | coint | Cointegration | Miscellaneous |
| 172 | coint_aeg_tstat_trend-ct_autolag-aic_maxlag-10 | coint | Cointegration | Miscellaneous |
| 173 | coint_aeg_tstat_trend-ct_autolag-bic_maxlag-10 | coint | Cointegration | Miscellaneous |
| 174 | pec | pec | Power envelope correlation | Miscellaneous |
| 175 | pec_orth | pec | Power envelope correlation | Miscellaneous |
| 176 | pec_log | pec | Power envelope correlation | Miscellaneous |
| 177 | pec_orth_log | pec | Power envelope correlation | Miscellaneous |
| 178 | pec_orth_abs | pec | Power envelope correlation | Miscellaneous |
| 179 | pec_orth_log_abs | pec | Power envelope correlation | Miscellaneous |

**TABLE S3:**
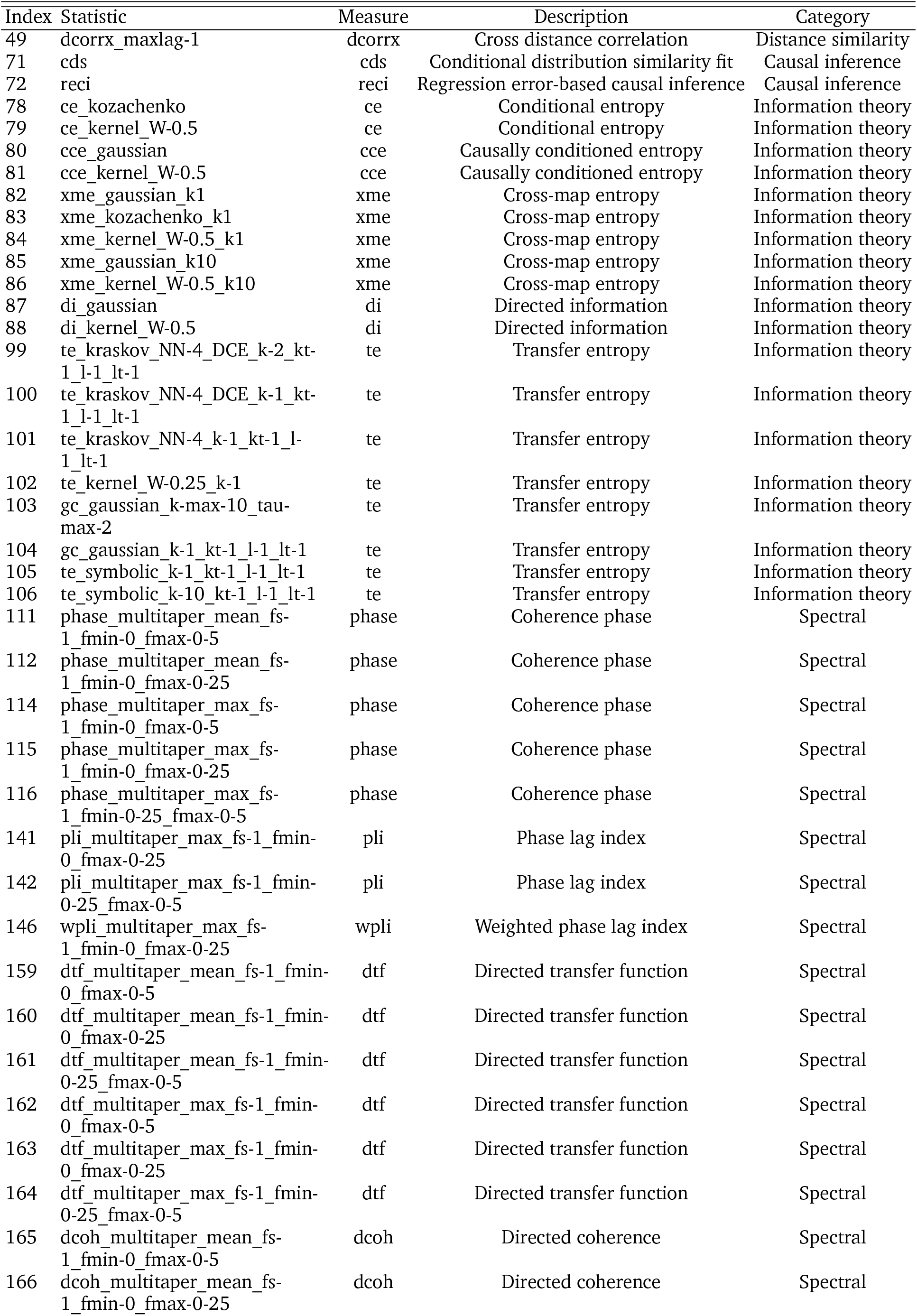
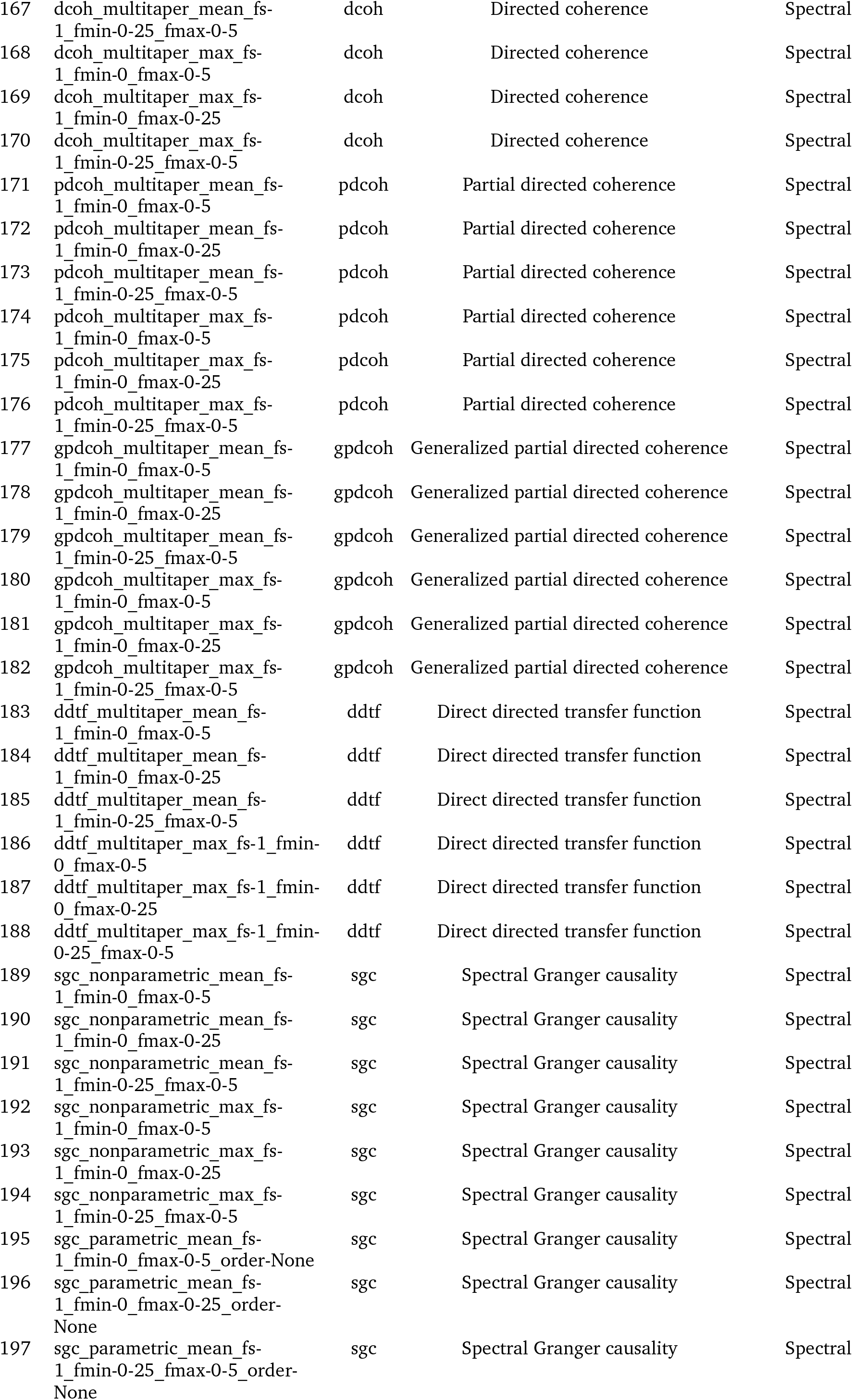
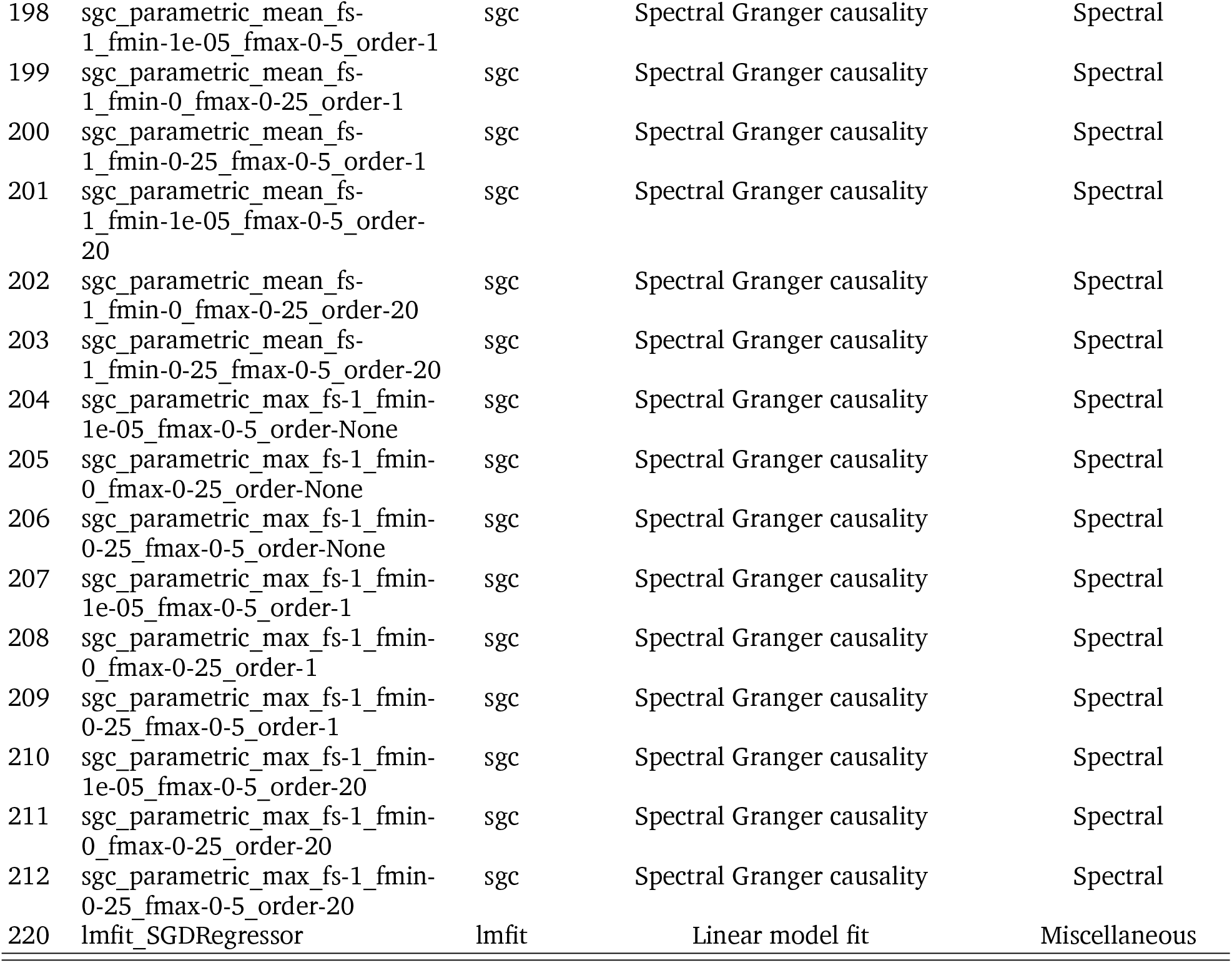
Directed *pyspi* statistics (excluding those only with opposite sign) Pairwise statistics with different upper and lower triangular values excluding those only differ by a sign.

**TABLE S4:** Directed *pyspi* statistics with opposite sign. Pairwise statistics with different upper and lower triangular values that only differ by a sign.

| Index | Statistic | Measure | Description | Category |
| --- | --- | --- | --- | --- |
| 73 | igci | igci | Information-geometric causal inference | Causal inference |
| 113 | phase_multitaper_mean_fs-1_fmin-0-25_fmax-0-5 | phase | Coherence phase | Spectral |
| 129 | psi_multitaper_mean_fs-1_fmin-0_fmax-0-5 | psi | Phase slope index | Spectral |
| 130 | psi_multitaper_mean_fs-1_fmin-0_fmax-0-25 | psi | Phase slope index | Spectral |
| 131 | psi_multitaper_mean_fs-1_fmin-0-25_fmax-0-5 | psi | Phase slope index | Spectral |
| 138 | pli_multitaper_mean_fs-1_fmin-0_fmax-0-5 | pli | Phase lag index | Spectral |
| 139 | pli_multitaper_mean_fs-1_fmin-0_fmax-0-25 | pli | Phase lag index | Spectral |
| 140 | pli_multitaper_mean_fs-1_fmin-0-25_fmax-0-5 | pli | Phase lag index | Spectral |
| 143 | wpli_multitaper_mean_fs-1_fmin-0_fmax-0-5 | wpli | Weighted phase lag index | Spectral |
| 144 | wpli_multitaper_mean_fs-1_fmin-0_fmax-0-25 | wpli | Weighted phase lag index | Spectral |
| 145 | wpli_multitaper_mean_fs-1_fmin-0-25_fmax-0-5 | wpli | Weighted phase lag index | Spectral |

**TABLE S5:**
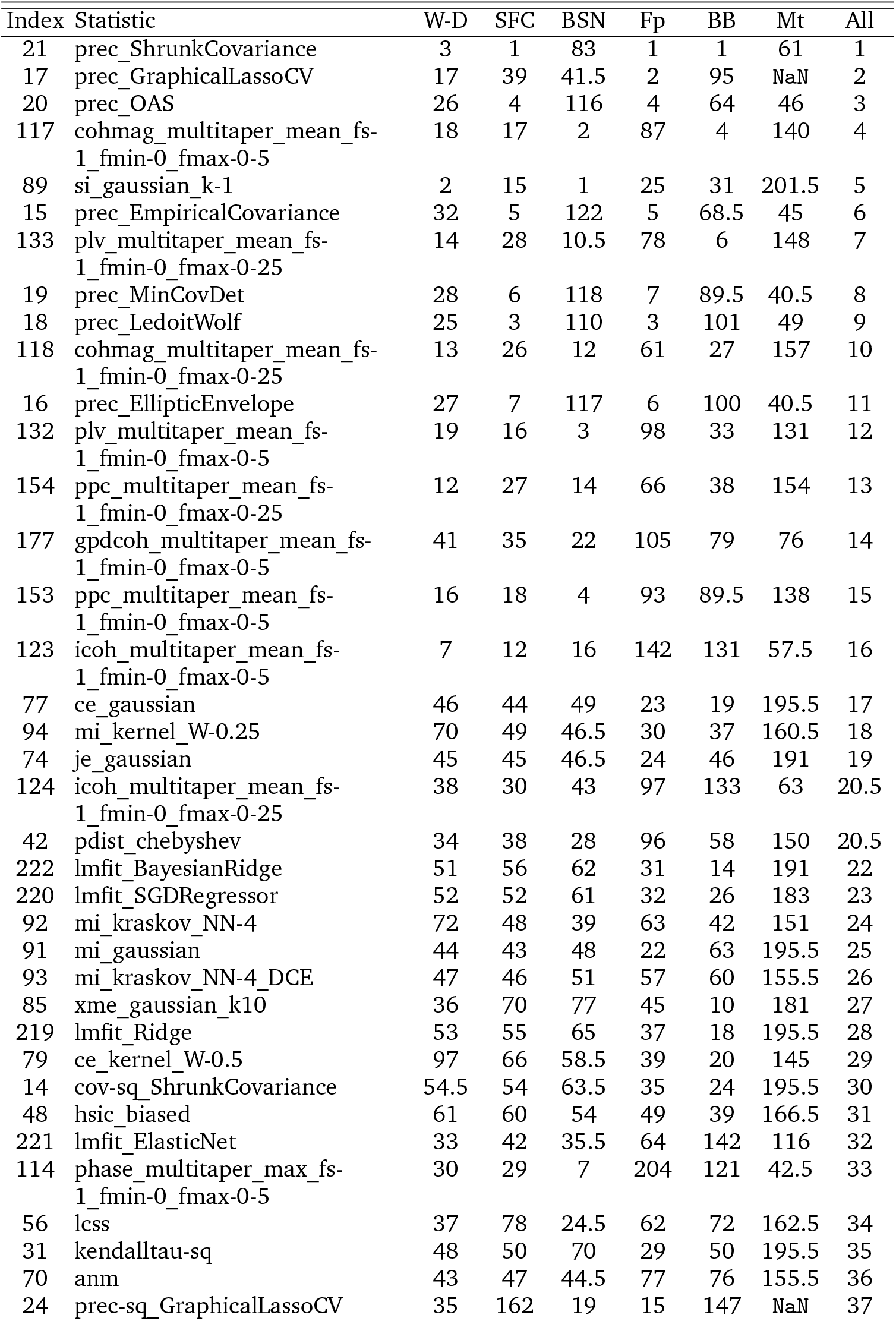
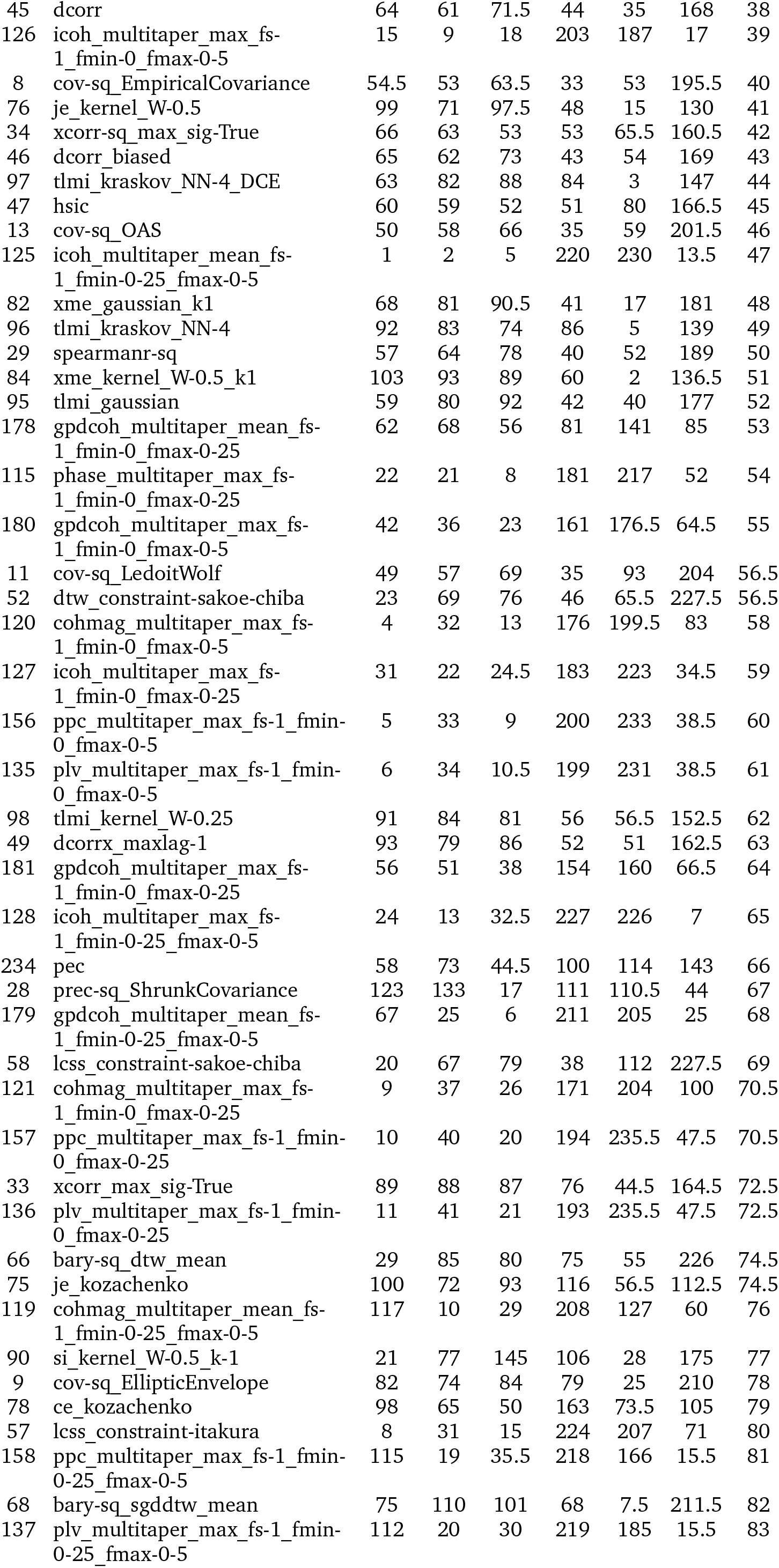
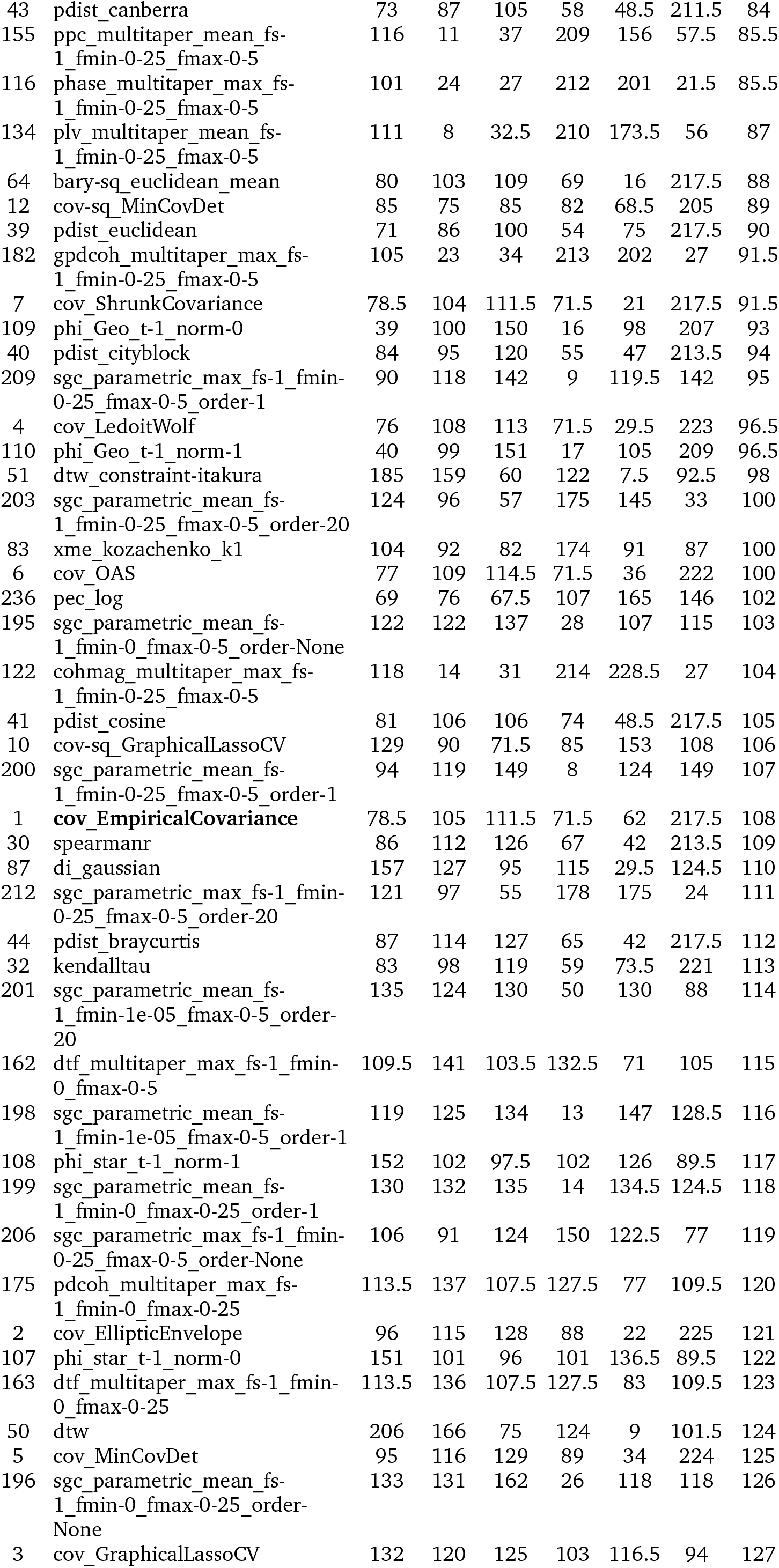
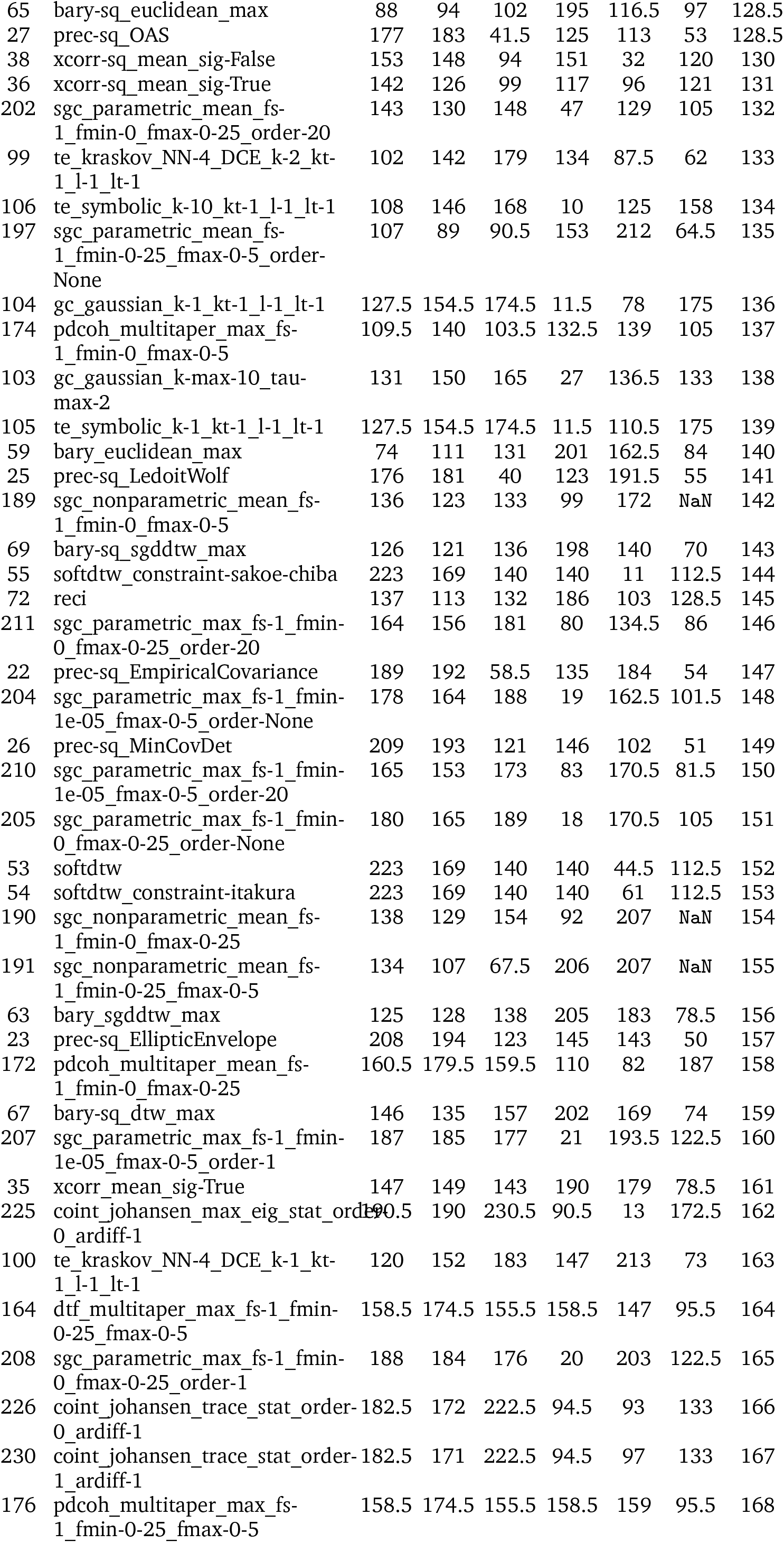
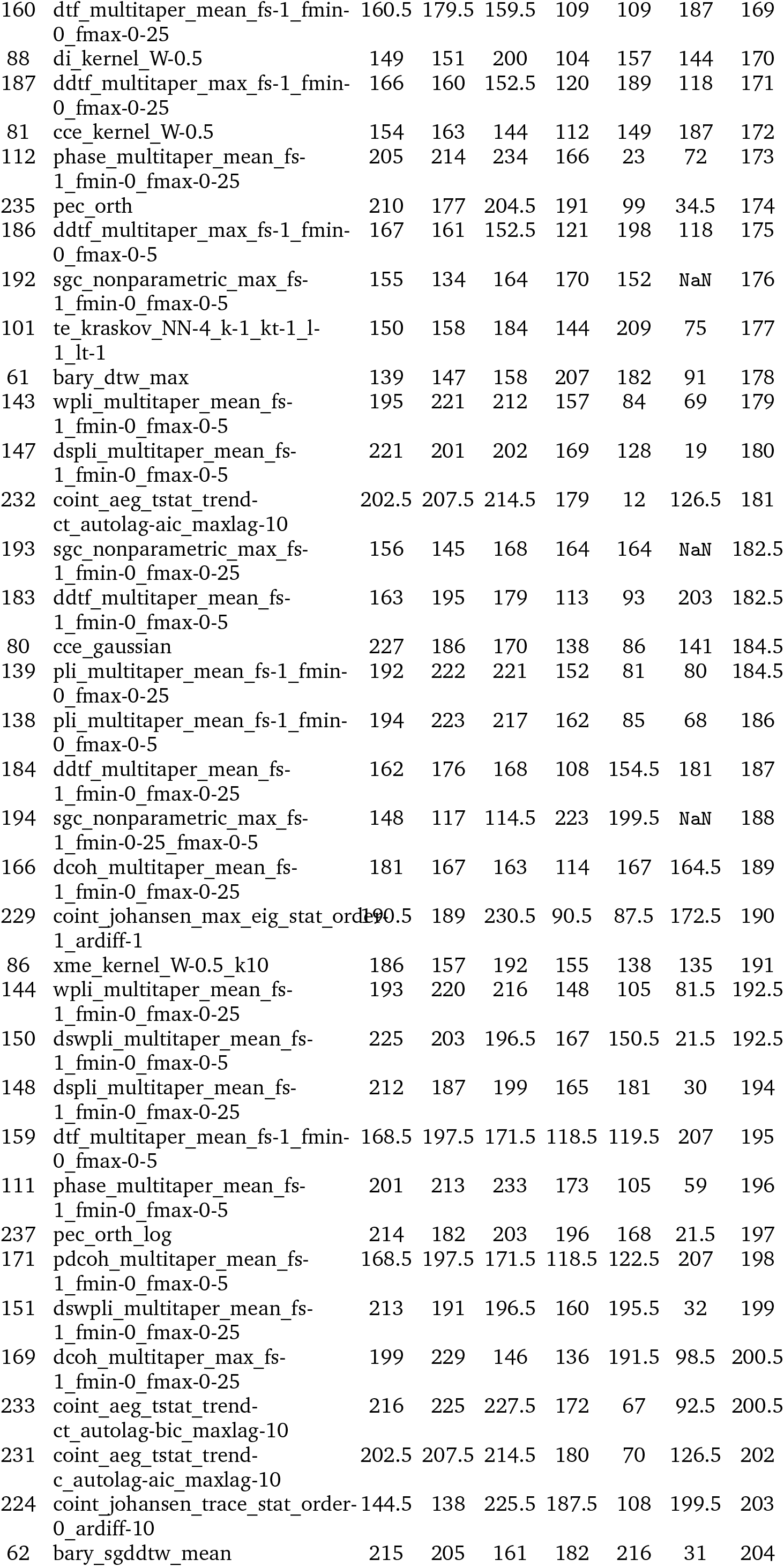
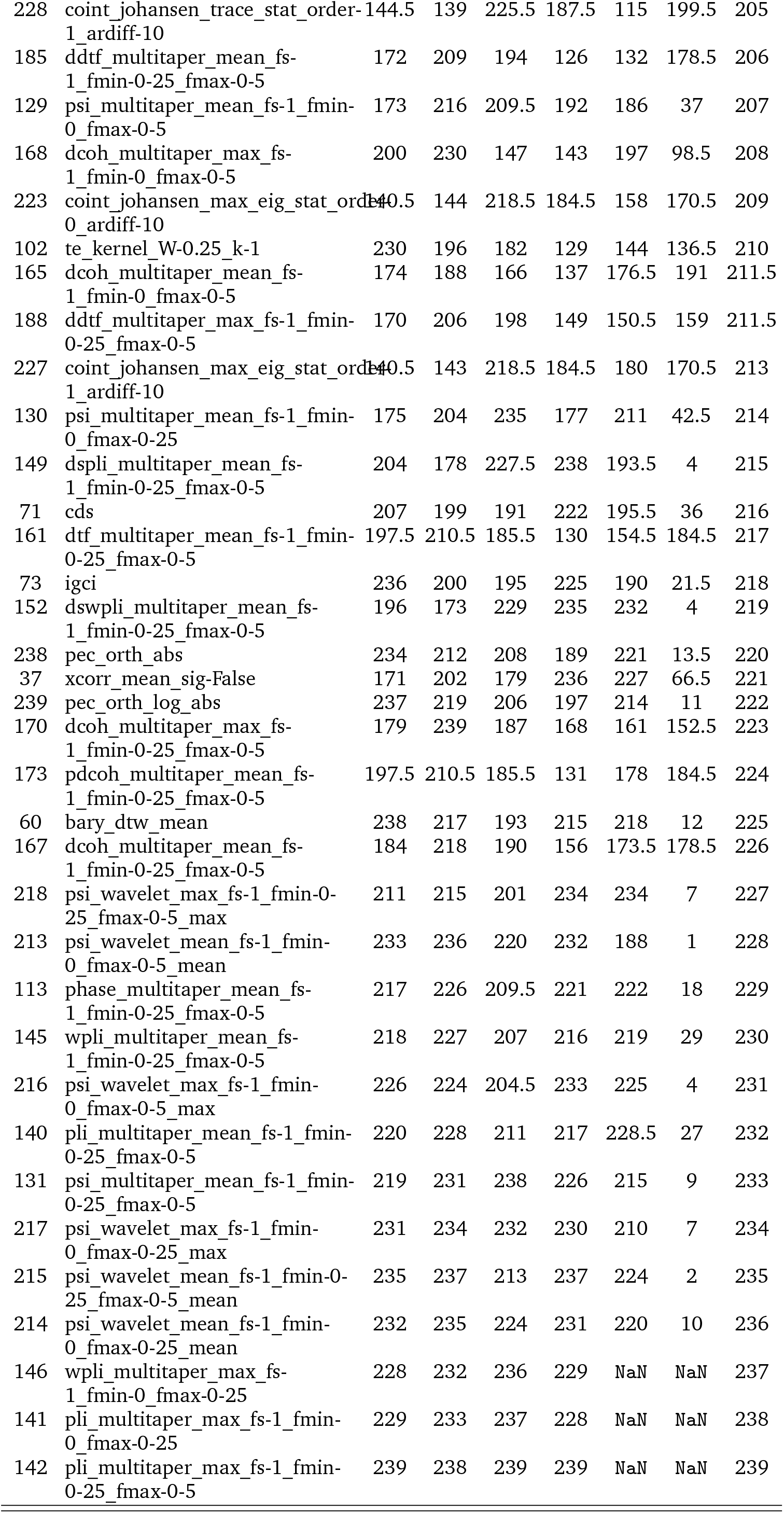
Individual and composite ranks of 239 *pyspi* statistics. Six potentially desirable criteria evaluated through the project were used to rank the 239 pairwise statistics. The criteria are: (1) negative weight–distance relationship (W-D), (2) positive structure–function coupling (SFC), (3) close correspondence with biological similarity networks (BSN), (4) high individual–participant identifiability (Fp), (5) high brain– behavior prediction (BB), and (6) low susceptibility to participant motion (Mt). Note that (3) was derived as a composite ranking averaging over the 5 neurophysiological networks in Fig. 3, (5) was derived as a composite ranking averaging over the 5 cognitive-behavior predictors in Fig. 4b, and (6) was derived as a composite ranking averaging over the two motion metrics in Fig. S11. The overall composite ranking (All) was derived by averaging the six individual rankings, and was used to reorder the final table. Tied elements were assigned ranks using the default strategy in *scipy*.*stats*.*rankdata*. NaN elements were ignored during the ranking.

